# Rewired mRNA homeostasis enforces cell size-associated senescence

**DOI:** 10.64898/2026.10.21.760624

**Authors:** Letizia Manfredi, Kaivalya Walavalkar, Shreyasi Neogi, Lisa Bucher, Falko Noé, Gabriel E Neurohr, Raffaella Santoro, Andrew S Belmont, Matthias Altmeyer, Sandhya Manohar

## Abstract

Cellular senescence refers to a permanent state of cell cycle exit and plays a key role in blocking the proliferation of compromised cells. Despite diverse drivers, senescent cells often share common features, one of which is their increased size. Though previously viewed as a passive outcome of cell growth during senescence, recent work shows that excess cell size can actively promote senescence via the p53-p21 pathway, making it both a consequence and a cause of cell cycle arrest. *How cells sense their own size and communicate this information to the cell cycle control apparatus remains unresolved.* Using genome-wide proximity mapping, we show that excess cell size globally redistributes chromatin between nuclear speckles and the nuclear lamina. These changes correspond with altered gene expression profiles, including the de-repression of typically silenced lamina-associated genes, some of which are involved in the senescence-associated secretory phenotype (SASP). Consistent with altered nuclear speckle function, we find that excess cell size perturbs normal splice isoform usage. In particular, *MDM4*, a negative regulator of p53 signaling, is differentially spliced when cells become enlarged to yield a non-functional variant, *MDM4-S*. Re-introducing correctly spliced *MDM4* transcripts restores G_1_ cell cycle progression, thereby reverting senescent phenotypes in enlarged cells. Together, these data indicate that excess cell size is sensed via changes in mRNA homeostasis associated with a reorganized genome, driving both the secretory phenotype and permanent cell cycle exit that are characteristic of cellular senescence.

## Introduction

Though the cells that comprise an organism span five orders of magnitude in size, cells of a given type typically occupy a narrow size range [1]. This narrow distribution is a consequence of cells systematically coupling their growth rates to division or differentiation [2–4] and suggests that cells have a rheostat-like mechanism for defining a size “set point.” Deviating from this set point negatively impacts cell physiology. Indeed, tumor cells display increased cell size heterogeneity [5], and this is associated with increased invasiveness [6]. Cells that are smaller than their set point undergo cell cycle delays in G_1_ and activate the stress responsive p38 kinase [7]. Cells that are too large also encounter problems: Excess cell size has long been associated with cellular senescence [8], and earlier work from our group and others has shown that it is also accompanied by an array of other cell cycle and genome homeostasis defects. Though many enlarged G_1_ cells are already senescent (likely a mechanism for preserving genome integrity), those that re-enter the cell cycle are prone to replication-stress, fail to repair DNA damage expediently, and tend to either skip or undergo erroneous mitoses [9–14].

Though it has been known since the 1960s that mammalian cells only initiate DNA synthesis after they have obtained a critical mass [15], how they sense their own size and communicate this to the cell cycle control apparatus remain open questions. Recent work has revealed that mammalian cell growth is intrinsically coupled to cell cycle entry via the dilution of the cell cycle inhibitor Rb as cells grow [16, 17]. While this provides a model for how cells sense when they are large enough to enter the cell division cycle, a corresponding mechanism has yet to be established for how cells sense that they are too large to faithfully execute the cell cycle. Earlier work from our group and others has shown that this involves a p53-dependent process [9–12], but it is not yet clear what signaling events converge upon p53 activation to restrain cell cycle progression in cells that are too large.

Here, we show that excess cell growth perturbs mRNA homeostasis at the gene expression and mRNA processing levels. Cellular enlargement redistributes chromatin between transcription-permissive nuclear speckles and the transcription-refractory nuclear lamina. This reorganization correlates with the de-repression of lamina-associated genes, some of which are involved in the senescence-associated secretory phenotype (SASP). We find that enlarged cells further alter their transcriptome by implementing size-dependent alternative splicing programs, possibly due to nuclear speckle dysfunction, resulting in widespread alterations in splice isoform usage. One such change occurs in a negative regulator of the p53 pathway, *MDM4*, which is alternatively spliced in enlarged cells to yield a non-functional variant, leading to unscheduled p53 activation that can be reversed by adding back correctly spliced *MDM4* cDNA. Together, these data suggest that cellular enlargement is sensed via transcriptional changes imparted by altered nuclear function, contributing to both senescence-associated gene expression and permanent cell cycle exit.

## Results

### Excess G_1_ cell size activates p53-p21 signaling

To study the effects of excess G_1_ cell size on cellular function, we used a previously established system combining reversible cell cycle arrests (mediated by the Cdk4/6 inhibitor palbociclib) with either growth restriction (mediated by co-treatment with the mTOR inhibitor Torin1) or temporally limited changes in arrest duration to modulate G_1_ cell size (**Fig 1A**) [9, 10, 12]. Using this strategy, cells that were arrested for 7 days significantly increased in volume relative to cycling cells, whereas cells that were co-treated with the mTOR inhibitor (mTORi) or arrested for a shorter duration (2d) were maintained close to physiological size (**Fig 1B**). Cells that were exposed to mTOR inhibition were co-treated for 6 days followed by an additional day of palbociclib alone to minimize pleiotropic effects of concurrent mTOR inhibition during downstream measurements. We hereafter refer to cells treated with palbociclib alone as “enlarged” G_1_ cells and cells that were co-treated with an mTOR inhibitor or treated for a shorter duration as “size-constrained” G_1_ cells. By focusing on phenotypes that were observed in enlarged cells relative to both methods of size-constraint, we identified changes that are size-dependent rather than solely a result of prior growth signaling restriction or differences in G_1_ arrest duration.

**Figure 1:**
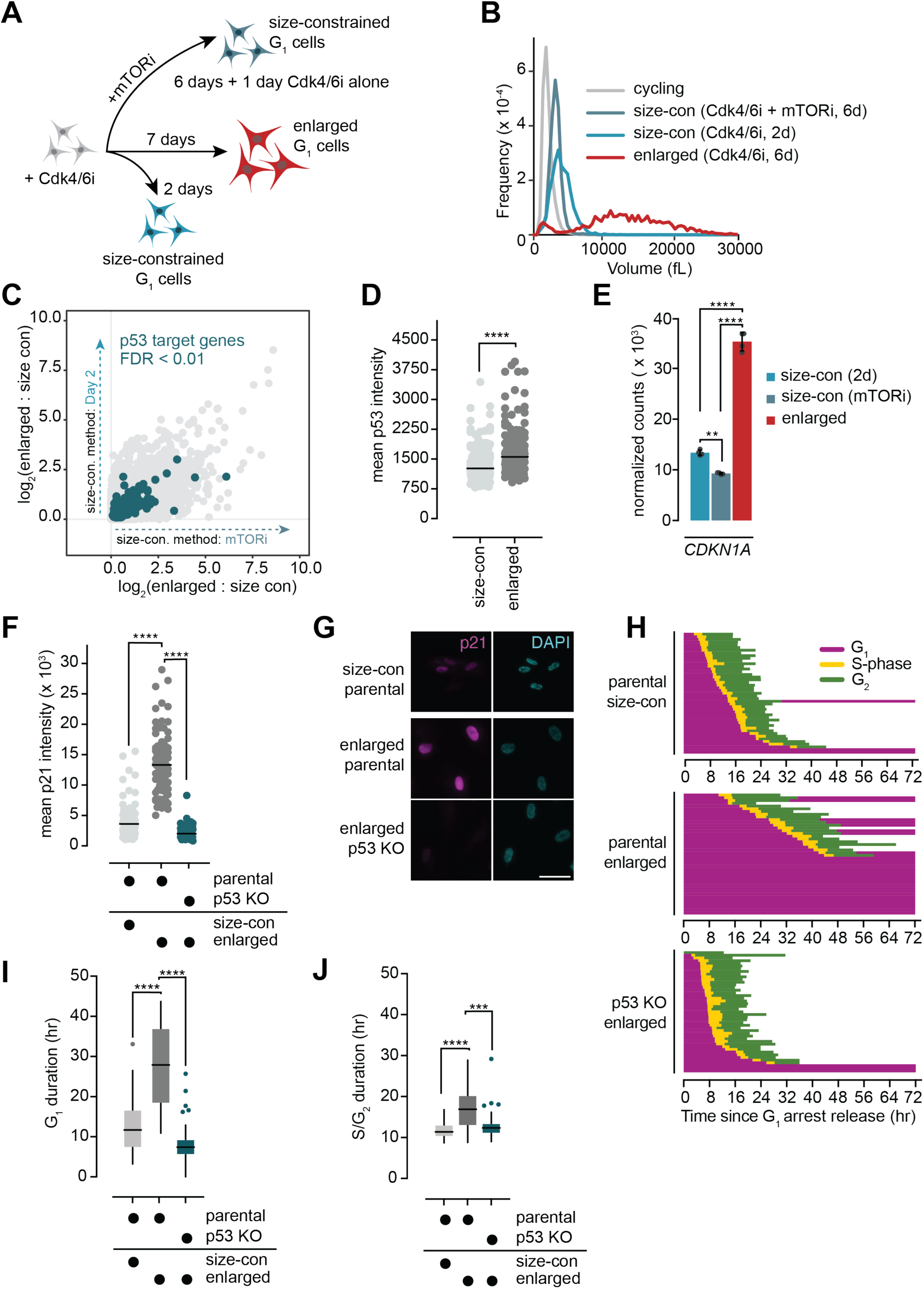
Excess G_1_ cell size activates p53-p21 signaling. **A.** Schematic for generating enlarged and size-constrained G_1_ RPE1 cells. **B.** Example cell volume measurements for cells treated as in (**A**). **C.** Comparison of upregulated transcript abundances in enlarged cells compared to each method of size-constraint (2d on the y-axis; mTORi on the x-axis). Annotated p53 target genes that are significantly upregulated (FDR < 0.01) are shown in blue, with 142 upregulated out of a target dataset of 342. **D.** Quantification of p53 mean nuclear intensity by immunofluorescence in enlarged and size-constrained (mTORi) cells. Bar indicates the median value. **** : p < 0.0001 by two-tailed, unpaired t-test. n ≥ 184 cells per condition. **E.** *CDKN1A* (p21) transcript abundance in enlarged and size-constrained cells as measured by RNA-seq. Data are reported as mean ± SD from four replicates. ** : p = 0.0014; **** : p < 0.0001 by one-way ANOVA followed by Tukey’s multiple comparisons test. **F.** Quantification of p21 mean nuclear intensity by immunofluorescence in enlarged and size-constrained (mTORi) cells. Measurements in enlarged cells were made in a parental and p53 knockout background. Bar indicates the median value. **** : p < 0.0001 by one-way ANOVA followed by Tukey’s multiple comparisons test. n ≥ 81 cells per condition. **G.** Example images of the experiment depicted in (**F**). Scale bar = 50 µm. **H.** Single cell traces of the size-constrained (mTORi) and enlarged RPE1 FUCCI cells released from a G_1_ arrest based on live cell imaging of FUCCI reporters. The end of a trajectory indicates mitotic entry or the end of the imaging duration. n = 45 cells for each condition. **I.** Tukey plots of G_1_ duration for cells that re-entered the cell cycle in the experiment depicted in (**H**). **J.** Tukey plots of S/G_2_ duration for the experiment depicted in (**H**). The end of S/G_2_ was determined by either mitotic entry or reversion to a G_1_ state based on the FUCCI markers.

Using the strategy outlined above, we and others previously demonstrated that enlarged G_1_ cells upregulate p53 signaling relative to size-constrained cells [9, 10]. To investigate this further, we performed whole-transcriptome sequencing to evaluate the impact of cellular enlargement on the expression of p53 target genes. We found that, out of the 342 annotated p53 target transcripts [18] identified in our dataset, more than 40% significantly increased (FDR < 0.01) in abundance in enlarged G_1_ cells (**Fig 1C**, **Fig S1A**). These findings correlated with an increase in nuclear p53 protein expression (**Fig 1D**) as well as increased expression of *CDKN1A*—which encodes p21, the key mediator of cell cycle arrest when p53 is active— at both the transcript (**Fig 1E**) and protein level (**Fig 1F-1G**) in enlarged cells. Consistent with a p53-dependent mechanism of p21 upregulation, the increase in p21 protein abundance we observed in enlarged cells was reversed under p53 knockout conditions (**Fig 1F-1G**).

Our previous work established that active p53-p21 signaling causes cell cycle progression defects in enlarged cells released from G_1_ [9]. These defects included G_1_ delays and arrests as well as mitotic skipping in enlarged G_2_ cells (“G_2_ slippage”). Using RPE1 FUCCI cells to track cell cycle progression [19], we confirmed these previous findings (**Fig 1H-1J**). Importantly, p53 deletion in enlarged cells reverted cell cycle progression to the pattern observed in size-constrained cells (**Fig 1H-1J**) [9–11], indicating that the delays, arrests, and defects observed in enlarged cells were a consequence of active p53 signaling.

The G_2_ slippage phenotype observed in enlarged cells that re-enter the cell cycle was characterized as a p53-dependent response to replication stress-induced DNA damage [9, 10, 20, 21]. Though the G_1_ delays and arrests we observed were also p53-dependent, it was not clear what activates p53 signaling in enlarged G_1_ cells prior to S-phase entry. We previously ruled out G_1_ DNA damage signaling as a potential cause for this: Enlarged G_1_ cells generated using our treatment scheme displayed no measurable DNA damage, and p21 induction still occurred if DNA damage signaling pathways were silenced [9]. Though p53 activation in enlarged cells may involve a p38-dependent mechanism [10], our transcriptomic (**Fig S1B**) and phosphoproteomic data [22] did not show a clear p38 induction signature (**Fig S1C**), and concurrent pharmacological inhibition of p38 during a palbociclib-mediated arrest constrained cell growth (**Fig S1D**), which could partially rescue cell cycle progression defects. We thus sought to investigate additional explanations for why excess G_1_ cell size could promote p53 signaling.

### Nuclear speckles subscale relative to cell size

Using the strategy outlined in **Fig 1A**, we had previously performed proteomic measurements in enlarged and size-constrained cells to understand how the proteome is remodeled with respect to increasing cell size [9]. Consistent with data from others [23], we found that enlarged cells are significantly depleted in proteins involved in mRNA processing and splicing as well as mRNA transport (**Fig 2A**). We confirmed this by showing that several mRNA processing proteins are depleted in enlarged cells by western blot, including: PTBP1 and hnRNPK (ribonucleoproteins involved in mRNA processing), U2AF2 (a splicing factor), and CCAR1 (a recently discovered splicing fidelity factor [24]) (**Fig S2A**). Consistent with our proteomic findings, imaging revealed that enlarged cells are depleted in Sm proteins, a ribonucleoprotein family that form the core of the spliceosome, relative to size-constrained cells (**Fig 2B-2C**). Transcript sequencing also revealed a downregulation of genes associated with the mRNA processing GO term in enlarged cells relative to cells subjected to either method of size-constraint (**Fig S2B**). Taken together, these data confirmed that mRNA processing and splicing machinery subscale at the gene expression and protein level in enlarged G_1_ cells.

**Figure 2:**
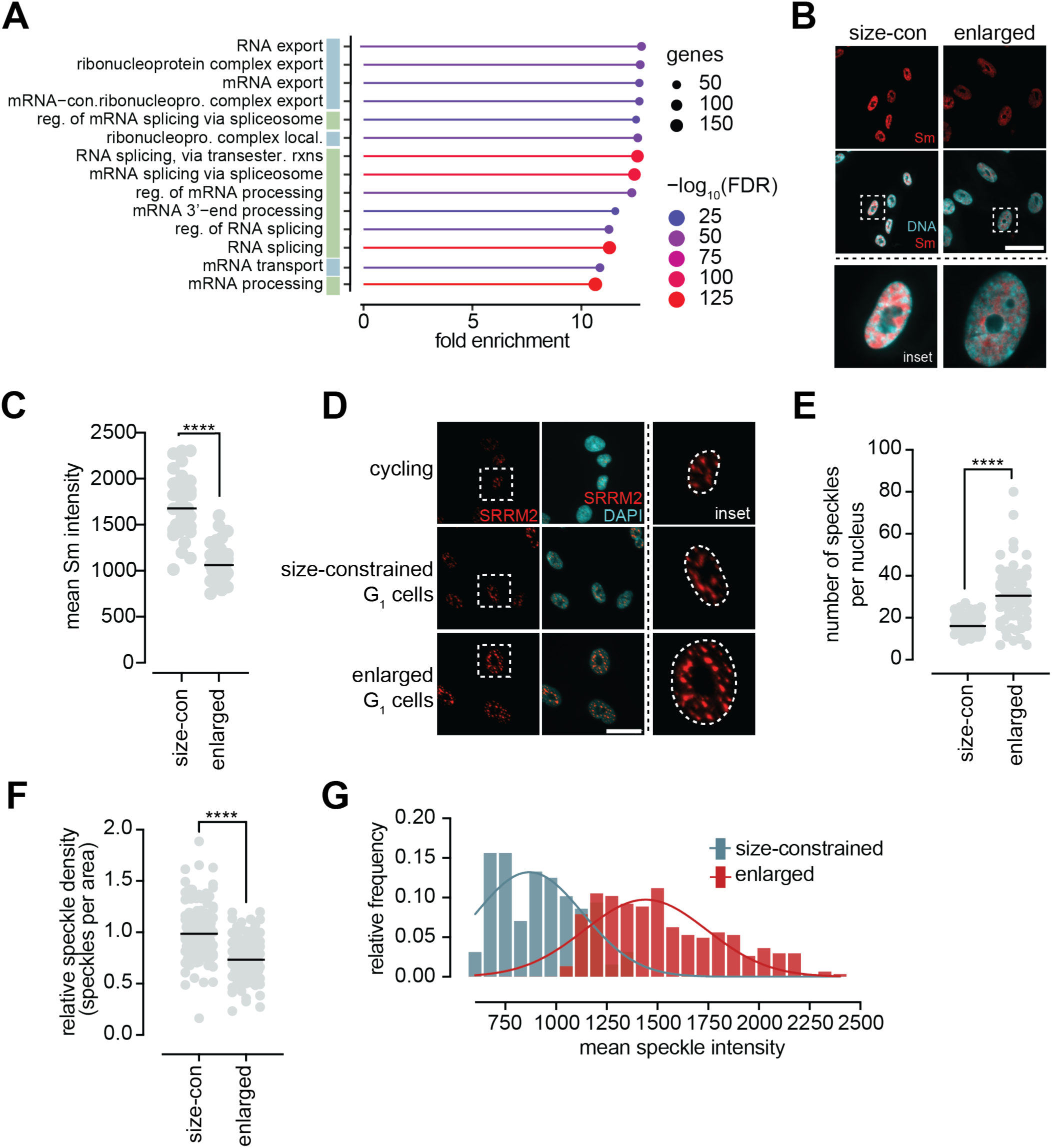
Nuclear speckles subscale relative to cell size. **A.** Gene ontology (GO) analysis of proteins that were significantly depleted in enlarged cells relative to both methods of size-constraint (2d and mTORi) in the proteomics dataset from Manohar, et al. *Mol Cell* (2023). Terms involved in mRNA transport are indicated in blue; terms involved in mRNA processing and splicing are indicated in green. **B.** Example images of enlarged and size-constrained (mTORi) cells that were stained for the Sm (Smith antigen) family of proteins. Scale bar = 50 µm. **C.** Quantification of the experiment shown in (B). Bar indicates median value. ****: p < 0.0001 by two-tailed, unpaired t-test. n = 38 cells per condition. **D.** Example images of cycling, enlarged, and size-constrained (mTORi) cells that were stained with the SC35 antibody directed against the speckle-resident protein SRRM2. Scale bar = 20 µm. **E.** Quantification of the number of speckles per nucleus for the experiment shown in (**D**). Bar indicates median value. **** : p < 0.0001 by two-tailed, unpaired t-test. n = 70 cells per condition. **F.** Speckles per nuclear area measured for the experiment shown in (**D**). Bar indicates median value. **** : p <0.0001 by two-tailed, unpaired t-test. n ≥ 106 per condition. **G.** Histogram of mean speckle intensity measured for the experiment shown in (**D**). n ≥ 128 per condition.

Many of the mRNA processing and splicing factors that were depleted in enlarged cells localize to nuclear speckles, which are sub-nuclear condensates that serve as hubs for mRNA maturation [25, 26]. Despite an overall downregulation of nuclear speckle-associated proteins, we found that the overall abundances of SRRM2 and SON, the two core components of nuclear speckles [27], are largely preserved in enlarged cells relative to size-constrained cells (**Fig S2C-S2D**). Immunostaining of SRRM2 (using an SC35 antibody [27]) confirmed that, although bulk nuclear SRRM2 concentrations were similar between enlarged and size-constrained cells (**Fig S2E**), their distribution, morphology, and intensity were altered. Though enlarged cells contained more nuclear speckles than size-constrained cells (**Fig 2D-2E**), they were less densely distributed within the nucleus (**Fig 2F**, **Fig S2F-S2G**). Moreover, whereas size-constrained cells had mostly small, irregularly shaped speckles, nuclear speckles in enlarged cells were generally enlarged and bright (**Fig 2D**, **Fig 2G**, **Fig S2G-S2H**). This phenocopied conditions of nuclear speckle dysfunction: Indeed, nuclear speckle enlargement has been shown to correlate with an accumulation of polyA(+) RNA within speckles [28], and increased speckle intensity, size, and circularity have been observed under conditions where splicing and/or transcription are compromised [29, 30].

Together, these observations indicated that nuclear speckles and many of their associated components subscale relative to nuclear volume in enlarged cells. This finding, coupled with the fact that enlarged G_1_ cells harbor the same DNA content as size-constrained G_1_ cells (**Fig S2I**) despite their increased nuclear volume (**Fig S2J**) [1, 3, 31, 32], made us consider whether the positioning of the genome relative to nuclear speckles may be altered as a consequence of cellular enlargement. Because genome organization around nuclear speckles can drive both gene expression [33–35] and mRNA splicing efficiency [36], this—along with alterations in mRNA processing factor abundance—could potentially compromise individual mRNA expression patterns in enlarged cells despite an overall scaling of bulk transcript abundance with cell size [37].

### Genome organization around nuclear speckles is altered as a function of increased cell size

In order to test the hypothesis that cellular enlargement perturbs normal nuclear speckle-genome contacts, we performed Tyramide Signal Amplification sequencing (TSA-seq), a genome mapping technique that measures the association between chromosomal loci and subnuclear compartments using an antibody coupled to tyramide free-radical based biotinylation of proximal DNA (**Fig 3A**). To map changes in nuclear speckle-genome interactions in enlarged cells, we performed TSA-seq using a SON antibody (**Fig S3A-S3C**), as has been done previously [29, 33–35].

**Figure 3:**
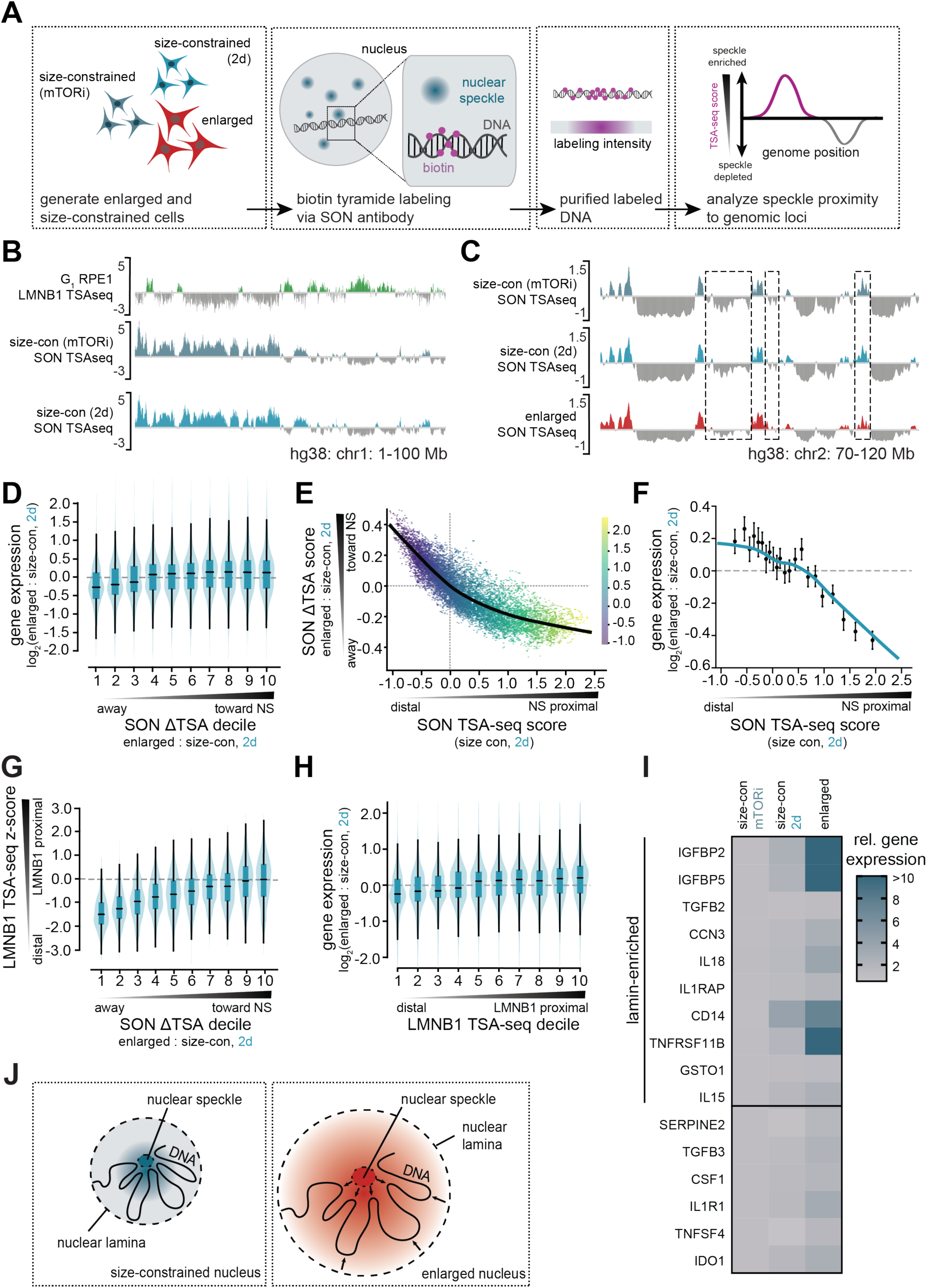
Genome organization around nuclear speckles is altered as a function of increased cell size. **A.** Schematic for the SON tyramide signal amplification (TSA) sequencing experiment. **B.** Genome browser tracks for the indicated TSA-seq experiments. LMNB1 TSAseq data were taken from Tran, et al. *Nucleic Acids Res* (2022), and SON TSAseq data were generated in this paper. The y-axis for all plots indicates z-score. **C.** Genome browser tracks for the indicated SON TSA-seq conditions. The y-axis for all plots indicates the TSA-seq score. **D.** Violin plots depicting gene expression for significantly changing (FDR < 0.05) genes in each SON ΔTSA decile for enlarged vs. size-constrained cells (2d). Spearman’s ⍴ = 0.18 between SON ΔTSA deciles and the indicated log_2_(fold change). n = 10,010 genes. **E.** Scatter plot depicting the change in SON ΔTSA score between enlarged and size-constrained cells (2d) as a function of the SON TSA-seq score in size-constrained cells (2d) for each gene. Spearman’s ⍴ = - 0.86. Color scale indicates the SON TSA-seq score in size-constrained cells (2d), and the black line depicts the LOWESS fit. n = 14,621 genes. **F.** Gene expression in significantly changing (FDR < 0.05) genes between enlarged and size-constrained cells (2d) plotted as a function of the SON TSA-seq score in size-constrained cells (2d). Spearman’s ⍴ = −0.24. Individual data points indicate the mean of 22 equal-sized gene bins with error bars indicating the 95% confidence interval. Blue line depicts the LOWESS fit. n = 10,010 genes. **G.** Violin plots depicting the Lamin B1 TSA-seq z-score for each SON ΔTSA decile for enlarged vs. size-constrained cells (2d). Spearman’s ⍴ = 0.46. n = 12,294 genes. **H.** Violin plots depicting gene expression changes in enlarged cells vs. size-constrained cells (2d) for significantly changing (FDR < 0.05) genes in each LMNB1 TSA-seq decile. Spearman’s ⍴ = 0.19. n = 8,592 genes. **I.** Gene expression changes (relative to mTORi) in the indicated SASP components that reposition closer to speckles in enlarged cells (SON ΔTSA_enl:size-con_ > 0 for both methods of size-constraint). Genes that are initially lamin-associated, based on the LMNB1 TSA-seq dataset, are highlighted. **J.** Graphical depiction of gene repositioning trends upon cellular enlargement.

The Pearson correlations between SON TSA-seq biological replicates were between 0.990 and 0.997, indicating a high degree of correlation (**Fig S3D**). Moreover, replicates from the two size-constrained conditions correlated more closely with each other than with enlarged cells. Principal component analysis corroborated this finding, indicating that the large majority of variance in our dataset is cell size-dependent, with only a small contribution from experimental differences (**Fig S3E**). Together, these results underscore that there was little experimental variability between biological replicates and that both size-constrained conditions represented similar biological states in terms of genome organization around nuclear speckles.

Nuclear speckle-associated domains (SPADs) are largely mutually exclusive from lamina-associated domains (LADs) [38, 39]. Comparing our data to a previously published lamin B1 chromatin TSA-seq dataset from G_1_ RPE1 cells [40], we found that speckle peaks were typically within inter-lamina-associated-domains (iLADs) whereas speckle valleys often corresponded to LADs, as expected. An example genomic region is shown in **Fig 3B**. Together, these observations underpinned that: (1) our SON TSA-seq measurements likely identified nuclear-speckle relationships accurately and that (2) the two methods of size-constraint employed in this study preserved normal chromatin architecture relative to untreated G_1_ RPE1 cells.

We found many regions wherein the magnitude of speckle-enrichment or speckle-depletion changed upon cellular enlargement (example region in **Fig 3C**). To quantify these changes at individual gene loci, we established the ΔTSA score (the difference in TSA-seq scores between two conditions) to measure the relative repositioning of loci between conditions. Consequently, a negative ΔTSA score indicates that a locus was more speckle-distal in a condition relative to a control, whereas a positive score indicates that it was more speckle-proximal. We observed a narrower distribution of ΔTSA values when comparing size-constrained cells to each other relative to enlarged cells vs. size-constrained cells (**Fig S4A**), suggesting that most of the repositioning we observed in our dataset is size-dependent.

There is a tight relationship between speckle proximity and gene expression levels, and even small positional changes can alter gene expression [34, 35, 41]. Based on this, we asked whether some of the gene expression changes associated with cellular enlargement could be correlated with altered speckle proximity. To investigate this, we binned the SON ΔTSA score for enlarged cells vs. both methods of size constraint into deciles and found that lower SON ΔTSA deciles correlated with reduced gene expression, whereas higher SON ΔTSA deciles correlated with increased gene expression (**Fig 3D**, **Fig S4B**). Together, these data indicated that the enlargement-associated repositioning of loci closer to nuclear speckles corresponded with increased gene expression, whereas gene repositioning further from speckles was associated with reduced gene expression.

We found that the genes with the highest SON TSA-seq scores (e.g., the most speckle proximal) tended to have the lowest SON ΔTSA score when comparing enlarged to size-constrained cells using either method of size-constraint (**Fig 3E**, **Fig S4C**). This indicated that speckle-proximal genes tended to reposition away from speckles upon cellular enlargement. This repositioning correlated with a significant decrease in gene expression, with the strongest gene expression response occurring at the most speckle-proximal loci (**Fig 3F**, **Fig S4D**).

In contrast, we found that lamina-associated genes tended to reposition towards nuclear speckles upon cellular enlargement (**Fig 3G**, **Fig S4E**). LADs typically represent a repressive chromatin environment and are characterized by low levels of gene expression [42]. We found that movement of lamina-associated genes towards speckles upon cellular enlargement was associated with the de-repression of these genes (**Fig 3H**, **Fig S4F**). Of note, we identified several upregulated senescence-associated secretory phenotype (SASP) factors [43–50], many of which are initially lamina-associated, that are re-positioned towards nuclear speckles in enlarged cells (**Fig 3I**).

Consistent with speckle-associated genes (which are typically early-replicating) repositioning away from speckles, we found that genes that were more speckle-distal upon enlargement tended to be annotated as early-replicating genes, whereas genes that were more speckle-proximal upon enlargement were late-replicating (**Fig S4G**). This is in agreement with lamina-associated genes moving closer to speckles, since these are typically later-replicating than the genome as a whole [51].

In summary, cellular enlargement appears to reposition speckle-associated genes further from speckles and lamina-associated genes towards speckles (**Fig 3J**). In both cases, this repositioning is broadly correlated with decreased and increased gene expression, respectively. We identified several upregulated SASP factors that were repositioned towards nuclear speckles in enlarged cells, suggesting that some senescence-associated gene expression profiles in enlarged cells may be guided by chromatin repositioning between the nuclear lamina and nuclear speckles.

### Excess G_1_ cell size drives differential mRNA splicing patterns

Based on recent findings that nuclear speckle proximity influences splicing efficiency [30, 36] and our observation that enlarged cells were depleted in mRNA splicing machinery (**Fig 2A**, **Fig S2A-S2B**), we hypothesized that enlarged cells may have perturbed mRNA splicing compared to size-constrained cells. To test this hypothesis, we engineered an RPE1 cell line stably expressing a previously-reported bichromatic fluorescent reporter of alternative splicing [52]. This reporter relies on the expression of two mutually exclusive open reading frames that can be toggled between based upon the inclusion or exclusion of an exon: Its inclusion results in GFP expression, whereas its exclusion shifts the reading frame, resulting in dsRED expression (**Fig 4A**). We validated this reporter by silencing *SRSF2* (a splicing factor whose knockdown globally affects RNA splicing patterns [53]) and confirmed that this causes an accumulation of dsRED relative to GFP (**Fig S5A-S5B**). Using our splicing reporter, we asked whether there was a shift in exon inclusion vs. exclusion events in enlarged vs. size-constrained cells (**Fig S5C**). We found that enlarged cells expressed more dsRED than cycling or size-constrained cells, indicating greater expression of the exon-skipped construct, and a lower ratio of GFP/dsRED, indicating a shift in the balance between exon inclusion and exclusion toward exon exclusion (**Fig 4B-4C**). Together, these data supported our hypothesis that cellular enlargement perturbs normal mRNA splicing.

**Figure 4:**
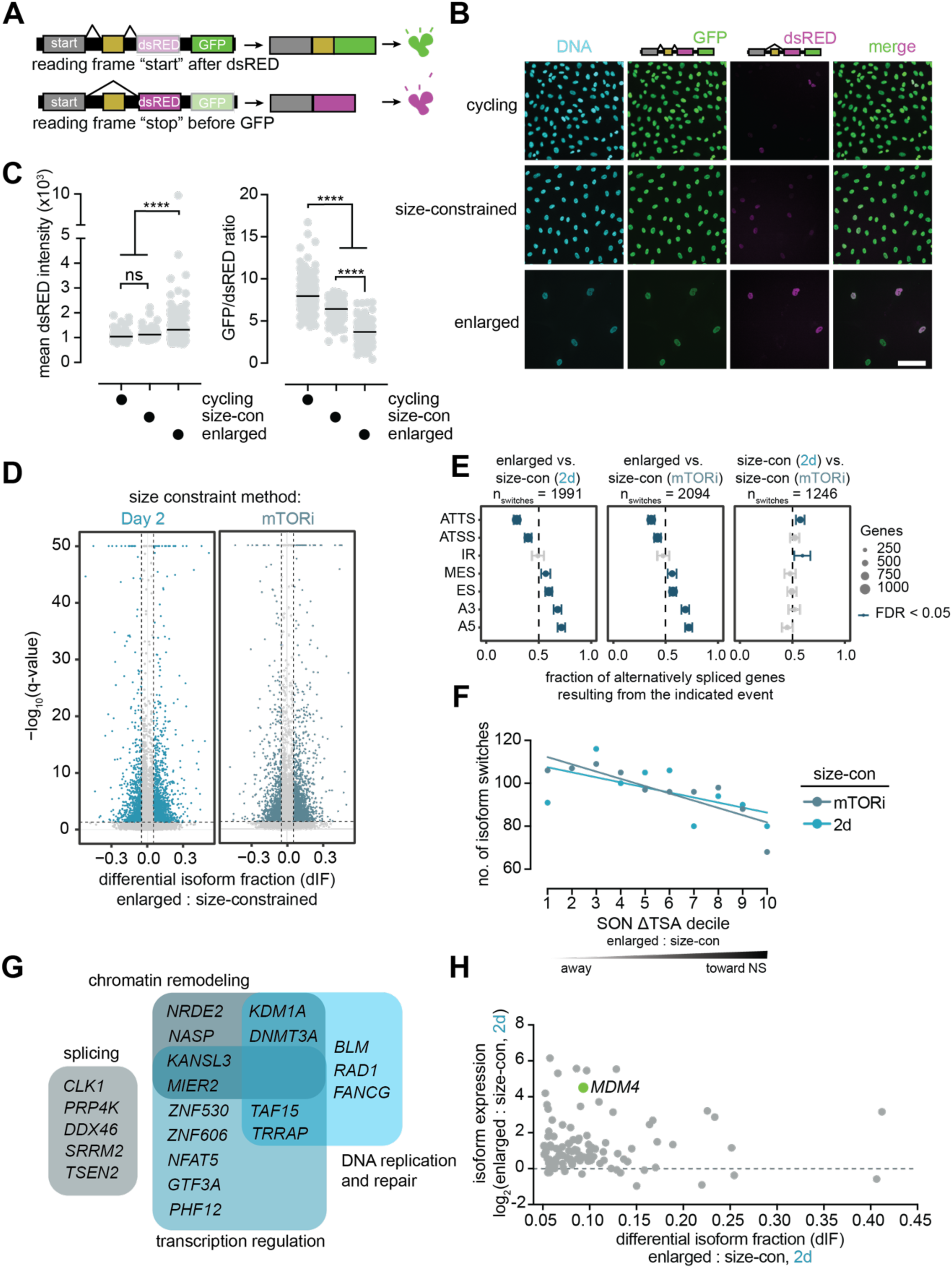
Excess G_1_ cell size drives differential mRNA splicing patterns. **A.** Schematic for the bichromatic splicing reporter used in (**B**) and (**C**). **B.** Example images of cycling, size-constrained (2d), and enlarged cells stably expressing the RG6 splicing reporter construct. Scale bar = 100 µm. **C.** Quantification of the mean dsRED intensity (left) and GFP/dsRED ratio (right) per cell in the experiment shown in (B). Bar indicates median value. n ≥ 170 cells per condition. **** : p < 0.0001; ns : p = 0.091 by one-way ANOVA followed by Tukey’s multiple comparisons test. **D.** Volcano plots depicting the differential isoform fraction (the relative proportion of a specific transcript isoform coming from a single parent gene) for a given transcript on the x-axis relative to the isoform switch q-value (y-axis) for enlarged vs. size-constrained cells (right, mTORi; left, 2d). Colored data points represent those with q < 0.05 and a dIF > 0.05. **E.** Enrichment of alternative splicing events among genes with isoform switches for the indicated comparisons. Data points depict the fraction of genes in which the isoform gaining usage in enlarged cells (left and center panel) or size-con (2d) cells (right panel) carries the indicated event. Error bars indicate 95% confidence interval, and point size corresponds to the number of genes. Blue data points indicate an FDR < 0.05 by two-sided binomial test with Benjamini–Hochberg correction. ATTS: alternative transcription termination site, ATSS: alternative transcription start site, IR: intron retention, MES: multiple exon skipping, ES: exon skipping, A3: alternative 3′ splice site and A5: alternative 5′ splice site. **F.** The number of isoform switches observed per SON ΔTSA decile when comparing enlarged to size-constrained cells. **G.** Select PTC transcripts that are fractionally upregulated in enlarged cells, sorted by category. **H.** PTC transcripts that are fractionally upregulated (n = 104) in enlarged cells relative to size-constrained cells (2d), plotted as PTC isoform expression change vs. differential isoform fraction. Note that a PTC transcript can still have an increased dIF (fractional abundance) despite its expression level decreasing (overall relative abundance).

To investigate this hypothesis further on an endogenous, gene-specific level, we evaluated differential fractional splice isoform usage in our transcriptomic dataset (which contained ∼170,500 isoforms of ∼21,000 coding genes) (**Fig 4D**, **Fig S5D**). Pairwise comparisons of our three conditions revealed a systematic bias towards particular alternative splicing events in enlarged vs. size-constrained cells but not when comparing the two methods of size-constraint to each other (**Fig 4E**). Most notably, we found that enlarged cells tended to undergo multiple- and single-exon skipping events and alternative 3’- or 5’-splice site usage relative to size-constrained cells. Together, these data indicated that there was an increased frequency of alternative transcript isoform usage in enlarged cells relative to size-constrained cells, and these changes resulted from specific changes in mRNA processing.

Because the efficiency of co-transcriptional mRNA splicing is linked to gene-speckle proximity [36], we asked if there was a relationship between alternative splicing and gene repositioning relative to speckles in enlarged cells. Though we observed a general decrease in the number of alternative splicing events observed in genes in higher SON ΔTSA deciles (i.e., genes that reposition closer to nuclear speckles in enlarged cells are less frequently alternative spliced) (**Fig 4F**), we did not observe a systematic relationship between particular alternative splicing events and repositioning directionality relative to speckles (**Fig S5E-S5F**). Thus, global changes in speckle proximity were unlikely to explain the specific alternative splicing biases we observed in enlarged cells.

Though some alternative splicing events are inert, others can significantly impact transcript stability. Premature termination codons (PTCs), for example, result in nonsense-mediated mRNA decay (NMD), causing a loss of transcript expression. To identify extreme cases of alternative splicing events that could impact the expression of relevant genes, we focused on PTC transcripts with increased fractional usage in enlarged cells. We detected several DNA replication and repair transcripts that undergo isoform switching to yield PTC transcripts in enlarged cells (**Fig 4G**). This could potentially contribute to the subscaling of DDR factors [23] and the propensity for replication stress and DNA damage previously observed in enlarged cells [9]. PTC transcript induction also occurred in several genes involved in splicing, suggesting that splicing aberrations may be self-reinforcing upon cellular enlargement (**Fig 4G**). Lastly, we observed PTC isoform switches in chromatin remodelers and transcription regulators, potentially contributing to genome organization changes as well as transcriptional changes in enlarged cells (**Fig 4G**). Thus, size-dependent alternative splicing impacts diverse cellular pathways and could contribute to many of the defects that we and others have observed in enlarged cells. In our ranked list of induced PTC transcripts, one of the top hits we observed was *MDM4* (**Fig 4H**), a negative regulator of the p53-p21 pathway.

### MDM4, a negative regulator of p53, is alternatively spliced in enlarged G_1_ cells

MDM4 and MDM2 are two major negative regulators of the p53 pathway (**Fig 5A**), and both have been implicated in driving p53-mediated cell cycle exit as a consequence of alternative splicing in the absence of DNA damage [54–58]. Our transcriptome-wide analysis of gene expression patterns in enlarged vs. size-constrained cells revealed a significant increase in *MDM2* transcript abundance in enlarged cells (**Fig S6A**), likely due to an established feedback loop between p53 activity and *MDM2* expression [59, 60]. Nonetheless, this is not sufficient to fully suppress p53 signaling in enlarged cells. Moreover, we concluded that the increased *MDM2* expression we observed in enlarged cells is unlikely a consequence of size-dependent differential mRNA splicing since we observed minimal difference in its fractional isoform usage (**Fig S6B**). Of note, neither MDM2 nor MDM4 peptides were detected in our previous proteomic dataset [9].

**Figure 5:**
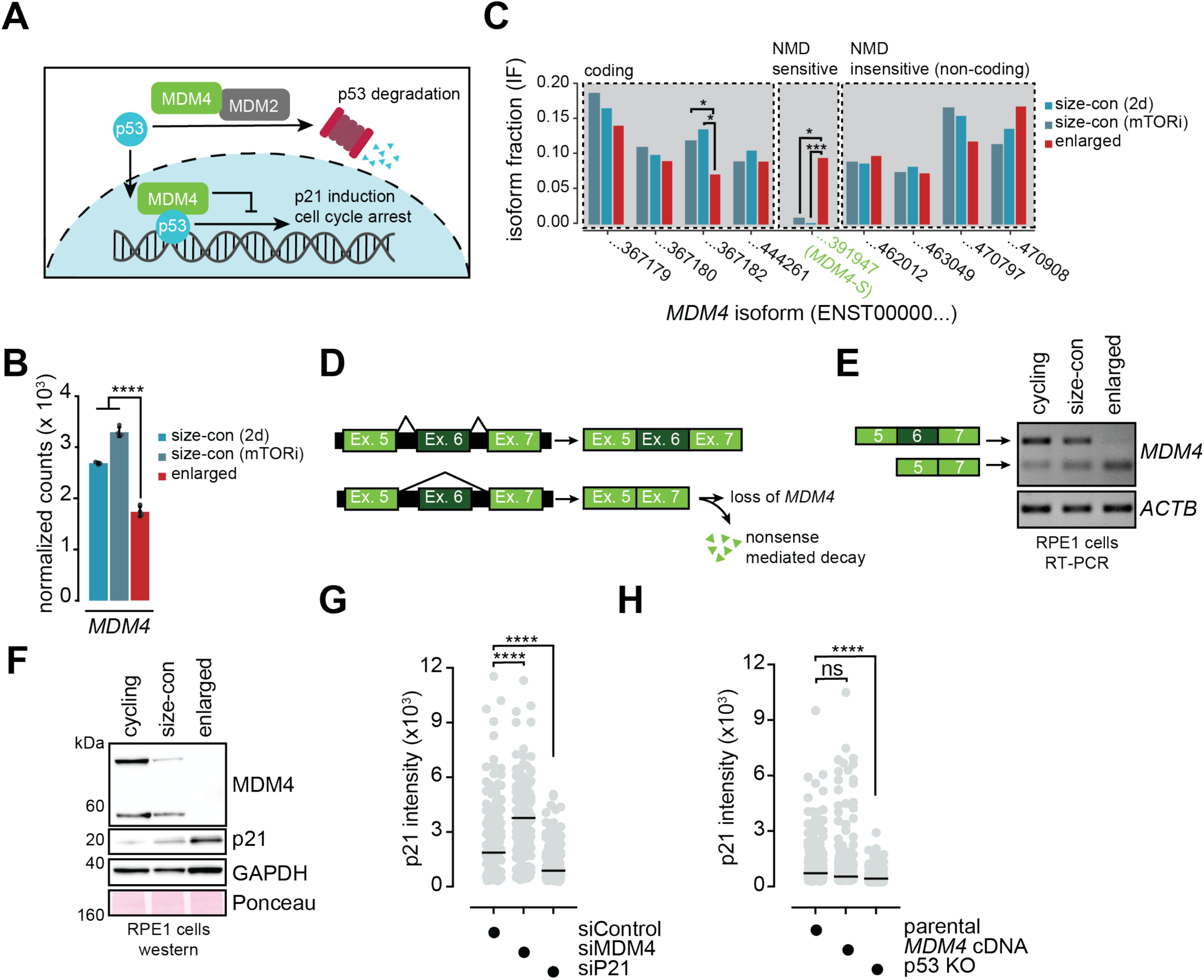
MDM4, a negative regulator of p53, is alternatively spliced in enlarged G_1_ cells. **A.** Schematic for the regulatory function of MDM2/MDM4 on p53 activity. **B.** *MDM4* total transcript abundance in enlarged and size-constrained cells, as measured by RNAseq. Data are reported as mean ± SD from four replicates. **** : p < 0.0001 by one-way ANOVA. **C.** Isoform fractions for the detected isoforms of *MDM4* in the RNAseq dataset in enlarged and size-constrained cells. *MDM4-S* is highlighted in green. * : 0.001 ≤ q < 0.05; *** : q < 0.001. **D.** Schematic for the Exon 5-6-7 junction in MDM4-FL (top) and MDM4-S (bottom). **E.** RT-PCR detecting *MDM4-FL* and *MDM4-S* in cycling, size-constrained (mTORi), and enlarged cells. ACTB was used as a loading control. **F.** Western blot indicating MDM4 and p21 protein levels in cycling, size-constrained (mTORi), and enlarged cells. GAPDH and Ponceau staining are used as loading controls. **G.** Quantification of mean p21 intensity by immunofluorescence in asynchronous RPE1 cells transfected with the indicated siRNAs. Bar indicates median value. **** : p < 0.0001 by one-way ANOVA followed by Tukey’s multiple comparisons test. n ≥ 150 cells per condition. **H.** Quantification of mean p21 intensity by immunofluorescence in the indicated asynchronous RPE1 cell lines. Bar indicates median value. **** : p < 0.0001; ns : p = 0.287 by one-way ANOVA followed by Tukey’s multiple comparisons test. n ≥ 313 cells per condition.

In contrast to *MDM2*, we found that *MDM4* total transcript abundance significantly decreases in enlarged cells relative to size-constrained cells (**Fig 5B**). On an isoform-specific level, we observed a >9-fold increase in the fractional usage of an NMD-sensitive isoform of *MDM4* called *MDM4-S* (which skips exon 6 and prematurely terminates in exon 7) in enlarged G_1_ cells (**Fig 5C-5D**). The *MDM4* locus did not appreciably reposition relative to nuclear speckles in our TSA-seq dataset (**Fig S6C**), although subtle positional effects or increased positional heterogeneity could not be excluded.

We validated our finding that enlarged cells accumulate the *MDM4-S* isoform by qRT-PCR (**Fig S6D**) and RT-PCR (**Fig 5E, Fig S6E**). Of note, because *MDM4-S* is subject to NMD, our steady-state measurements of this isoform likely underestimated its actual production in enlarged cells. Based on these findings, we hypothesized that alternative splicing of *MDM4* to yield high levels of the *MDM4-S* variant could result in a loss of functional MDM4 in enlarged cells, causing the unscheduled p53 pathway activation. Consistent with this, we observed a loss of MDM4 protein in enlarged cells relative to either size-constrained or cycling cells (**Fig 5F**).

To further test this hypothesis, we asked whether modulating MDM4 expression is sufficient to alter p21 abundance. We found that siRNA mediated knockdown of *MDM4* significantly increased both p53 and p21 levels (**Fig 5G**, **Fig S6F**) and blocked S-phase entry (**Fig S6G**). In contrast, however, overexpression of *MDM4* was not sufficient to decrease basal p21 levels (**Fig 5H, Fig S6H**). These data indicated that MDM4 is necessary but not sufficient to limit basal p53 signaling in RPE1 cells.

### Restoring full-length MDM4 transcripts in enlarged cells rescues cell cycle entry defects

To test our hypothesis that *MDM4* mis-splicing drives the p53 activity that causes cell cycle entry defects in enlarged cells, we asked whether restoring functional *MDM4* transcripts to enlarged cells is sufficient to rescue these defects. After confirming that long-term, ∼2-fold overexpression of *MDM4* cDNA did not constrain cell growth during a palbociclib-mediated G_1_ arrest (**Fig 6A, Fig S7A**), we measured p21 levels in parental and *MDM4* cDNA-expressing G_1_ cells that were either enlarged or size-constrained. We found that restoration of *MDM4* cDNA significantly reduced p21 levels in enlarged G_1_ cells but had no significant effect on basal p21 levels in size-constrained cells (**Fig 6B-6C**). As a control, we also measured p21 levels in enlarged and size-constrained p53 knockout cells, which ablated p21 expression in both cases (**Fig 6A-6B**).

**Figure 6:**
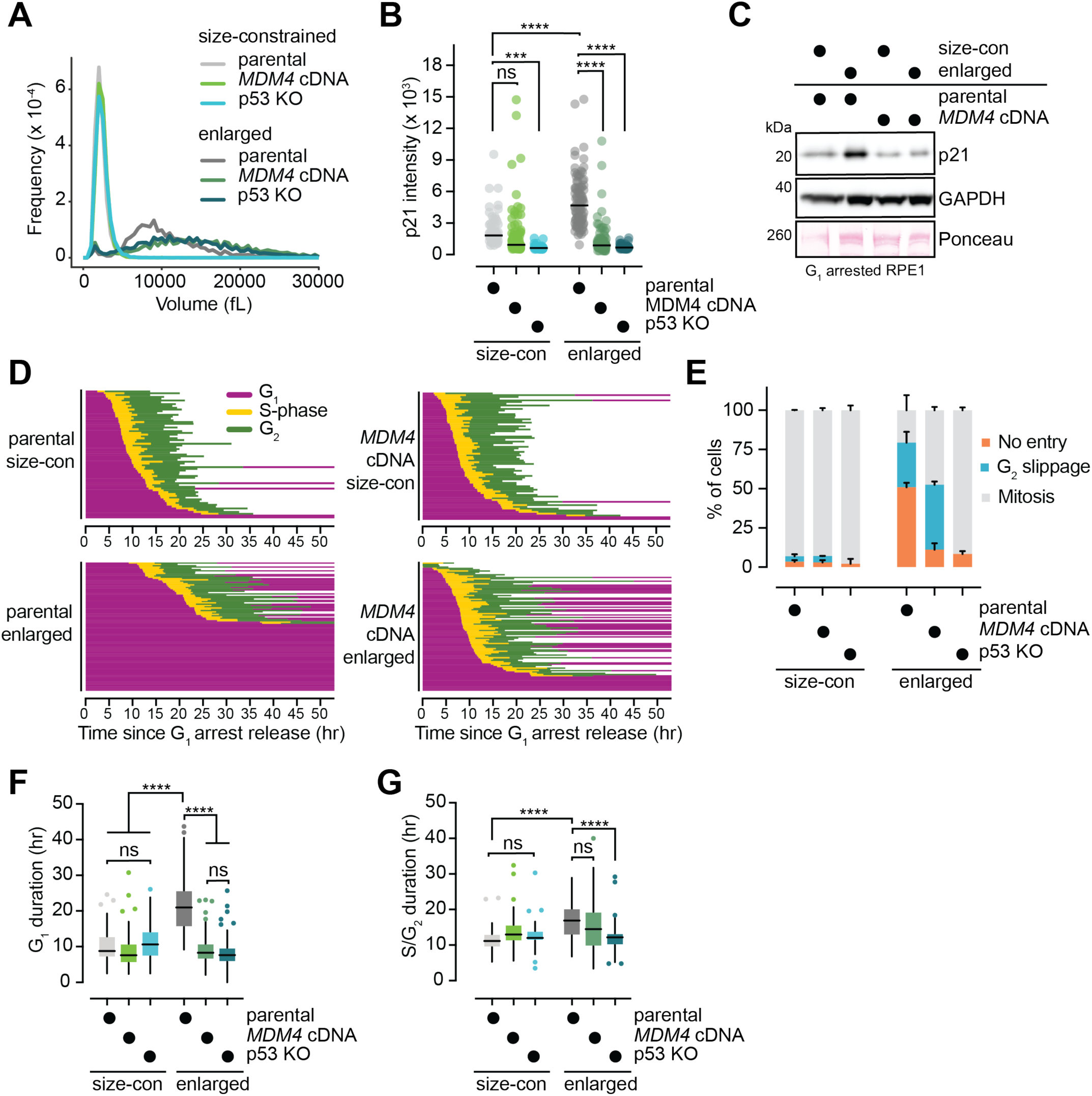
Restoring full-length MDM4 transcripts in enlarged cells rescues cell cycle entry defects. **A.** Example cell volume measurements for enlarged and size-constrained (mTORi) cells in the indicated RPE1 FUCCI stable cell lines. **B.** Quantification of p21 intensity by immunofluorescence in enlarged and size-constrained (mTORi) cells in the indicated RPE1 FUCCI stable cell lines. Bar indicates median value. n ≥ 68 cells per condition. **** : p < 0.0001; *** : p = 0.0008; ns : p >0.999 by two-way ANOVA followed by Šidák’s multiple comparisons test. **C.** Western blot indicating p21 protein levels in enlarged and size-constrained (mTORi) G_1_ cells in the indicated RPE1 FUCCI stable cell lines. GAPDH and Ponceau staining are used as loading controls. **D.** Single cell traces for size-constrained (mTORi) and enlarged cells in the indicated RPE1 FUCCI stable cell lines released from a G_1_ arrest based on live cell imaging of FUCCI reporters. The end of a trajectory indicates mitotic entry or the end of the imaging duration. n ≥ 70 cells per condition measured across 2-3 biological replicates. **E.** Quantification of cell cycle fates (no entry, G_2_ slippage, or mitotic entry) for the experiment shown in (**D**). Bars = mean with SD. **F.** Tukey plots quantifying G_1_ durations for cells that re-entered the cell cycle in the experiment shown in (**D**). **** : p < 0.0001; ns : p > 0.2 by one-way ANOVA followed by Tukey’s multiple comparisons test. **G.** Tukey plots quantifying S/G_2_ durations for the experiment shown in (**D**). **** : p < 0.0001; ns : p > 0.08 by one-way ANOVA followed by Tukey’s multiple comparisons test. The end of S/G_2_ was determined by either mitotic entry or reversion to a G_1_ state based on the FUCCI markers.

We next asked whether restoration of *MDM4* cDNA was sufficient to rescue the p53-dependent cell cycle defects we observed in enlarged cells. Using FUCCI cell cycle reporters, we found that restoring *MDM4* cDNA expression in enlarged cells rescued cell cycle entry defects, both by decreasing G_1_ delays in cells that re-entered the cell cycle and by eliminating the fraction of cells that were indefinitely arrested in G_1_ following release (**Fig 6D-6F**). In contrast, *MDM4* cDNA add-back had no significant effect on cell cycle progression in size-constrained cells (**Fig 6D-6G**). We orthogonally validated that restoring *MDM4* cDNA rescues S-phase entry in enlarged cells by measuring EdU incorporation following a G_1_ release in enlarged and size-constrained cells (**Fig S7B**). Consistent with its role in promoting cell cycle entry, we found that *MDM4* cDNA add-back improved long-term proliferation in enlarged G_1_ cells (**Fig S7C-S7D**). Importantly, we ruled out the possibility that high levels of *MDM4-S* (rather than a loss of *MDM4-FL*) drives the G_1_ cell cycle phenotypes observed in enlarged cells by demonstrating that excess *MDM4-S* expression was not sufficient to cause cell cycle defects in size-constrained cells (**Figure S7E**).

Though *MDM4-FL* cDNA restoration rescued cell cycle entry defects in enlarged cells, we found that this had no effect on the defects observed later in the cell cycle. Specifically, we found that restoration of *MDM4* cDNA did not rescue the fraction of enlarged cells subject to replication stress-dependent G_2_ slippage (**Fig 6D-6E**). Because G_2_ slippage is p53-dependent (**Fig 1H**), this finding indicated that restoring *MDM4* cDNA does not simply phenocopy p53 loss: It is sufficient to drive enlarged cells back into the cell cycle but is not sufficient to ablate p53 signaling in all contexts (for example, after replication-stress dependent DNA damage). Consistent with this, *MDM4* cDNA add-back was sufficient to alleviate the G_1_ delays in enlarged cells that re-entered the cell cycle (**Fig 6F**) but did not have a significant effect on replication stress-dependent S/G_2_ delays (**Fig 6G**). In contrast, p53 knockout rescued both G_1_ and S/G_2_ delays in enlarged cells (**Fig 1H-1J**; **Fig 6F-6G**). Supporting the idea that *MDM4* add-back does not unilaterally lower all p53 activity, we found that cells expressing *MDM4* cDNA still stabilize p53 following treatment with the DNA damaging agent doxorubicin (**Fig S7F**). These results suggested that MDM4 may not be sufficient to override p53 signaling under conditions that provoke DNA damage. In summary, restoring functional *MDM4* transcripts to enlarged G_1_ cells is sufficient to rescue cell cycle entry defects but not defects later in the cell cycle, suggesting that the effects we observed are not merely a consequence of over-expressing a negative regulator of p53 and are perhaps specific to the G_1_ defects observed in enlarged cells.

## Discussion

Here, we show that cellular enlargement redistributes chromatin between nuclear speckles and the nuclear lamina: Speckle-associated genes reposition away from nuclear speckles, and lamina-associated genes reposition towards nuclear speckles. While this repositioning could, in principle, be a consequence of the cellular flattening that is observed in senescent cells, our observation that this is accompanied by the de-repression of lamina-associated genes suggests a model in which lamina-associated genes exit LADs and migrate towards speckles in the nuclear interior rather than where the lamina is passively brought closer to speckles by flattening. This is consistent with the long-standing observation that senescent cells lose DNA-nuclear lamina contacts [61]. Our findings also agree with a recent study showing that nuclear speckles and the nuclear lamina compete to activate or repress some genes and that this balance can shift under conditions of speckle or lamina dysfunction [62].

We demonstrate that a subset of SASP-associated genes reposition closer to nuclear speckles, and this correlates with their increased expression. Interestingly, many of these are initially lamina-associated, and these are among the more highly upregulated repositioned SASP genes. Thus, some features of senescence-associated gene expression may be augmented by size-dependent genome reorganization. A similar chromatin repositioning pattern was recently observed during cortical neurogenesis: Lamina-associated genes in radial glia (many of which are involved in neuronal differentiation) re-localize to nuclear speckles upon differentiation into excitatory neurons, and this correlates with the increased expression of these genes [63]. Similar phenomena were observed during other forms of differentiation as well [64, 65]. Together, these and our findings raise the intriguing possibility that the chromatin repositioning can underpin cell fate decisions under circumstances that trigger changes in cell size, like differentiation or senescence, by influencing the associated gene expression profiles.

Additionally, we find that enlarged cells alter splice isoform usage patterns relative to size-constrained cells. Despite fewer alternative splicing events in genes that are repositioned closer to nuclear speckles in enlarged cells, we identified no systematic trends in particular alternative splicing events as a function of nuclear speckle proximity. In other words, the particular alternative splicing patterns we observe in enlarged cells cannot be clearly stratified by gene repositioning relative to nuclear speckles. This finding may instead be more broadly explained by the subscaling of splicing-associated factors as cells become larger [9, 23]. Why splicing machinery subscales relative to cell size remains an open question but could provide insight into why cellular enlargement, which entails corresponding nuclear enlargement [66], is tolerated without a fitness penalty under some physiological circumstances: Perhaps these cells are better able to scale their splicing machinery and thereby retain normal mRNA homeostatic programs. Supporting this possibility, splicing factors do not systematically subscale upon T-cell activation [67], which is associated with a 3-4-fold increase in cell volume [68].

Nonetheless, splicing aberrations appear to be a broad feature of senescent cells, occurring in replicative, stress-induced, and telomere-uncapping-induced senescence [55, 69, 70]. We found that *MDM4* mis-splicing is a feature of enlarged senescent cells that dictates their permanent cell cycle exit, and restoring correctly spliced *MDM4* rescues this. Though previous studies have linked Cdk4/6 inhibition to *MDM4* mis-splicing [71], our work clarifies that this is not a consequence of Cdk4/6 inhibition *per se* but rather a consequence of excess cell growth during a Cdk4/6 inhibitor-mediated arrest. Still, additional features of senescent cells have been attributed to other mis-splicing events: An exon skipping event in the translation regulator *EIF4H* was recently shown to promote the expression of some SASP genes in DNA damage-induced senescence [70], and the differential isoform fraction of *ING1* is altered during replicative senescence to remodel chromatin and modulate apoptosis susceptibility [72]. Together with these studies, our data suggest a model in which cells are exposed to an initial stress that causes cell cycle arrest, but continued cell growth introduces splicing aberrations that enforce various features of senescence to make it permanent. Further supporting this model, loss of splicing factors alone is sufficient to drive cells into senescence [55].

Lastly, our findings have important implications for addressing another yet-unanswered question about enlarged cells: Why are they prone to replication-stress? Our discovery that the genome is redistributed between typically early replicating SPADs and late replicating LADs could potentially play a role in enlarged cells’ tendency to undergo replication stress. Previous work has shown that the repositioning of LADs to the nuclear interior coincides with a shift from late to early replication timing [38, 73]. In the context of the large-scale rearrangement we observe upon cellular enlargement, this could potentially exhaust replication resources and alter the balance between replication, transcription, and chromatin assembly. Thus, our findings have the potential to explain diverse pathological features of enlarged, pre-senescent and senescent cells.

## Materials and Methods

### Cell culture, growth conditions, and drug treatments

hTERT-RPE1 cells and HEK293T cells (for lentivirus generation) were cultured in a humidified incubator at 37°C in the presence of 5% CO_2_. Wild-type hTERT-RPE1 cells were obtained from ATCC. hTERT-RPE1 FUCCI cells were a gift from Randall W. King with permission from the RIKEN Institute. All cell lines were regularly tested for mycoplasma contamination.

RPE1 cells were cultured in DMEM/F12 with GlutaMAX (Gibco) + 10% FBS and 1% penicillin/streptomycin. Cells that were allowed to grow large were seeded at ∼5,000-10,000 cells/cm^2^ and treated with 1-2 μM palbociclib (Sigma PZ0383 or Selleckchem S1116) for 7 days, maintaining sub-confluency throughout the experiment. Size-constrained cells that were generated using Torin1 (Sigma 475991) were seeded at ∼30,000 cells/cm^2^ and were cotreated with 1-2 μM palbociclib + 500 nM Torin1 for 6 days. After 6 days, cells were treated with palbociclib alone for an additional 24 hours to minimize the effects of Torin1 treatment in downstream experiments. For release experiments, cells were washed three times in media and released into media without drugs. HEK293T cells were cultured in DMEM with GlutaMAX (Gibco) + 10% FBS + 1% penicillin/streptomycin. Other drugs used in this study include: SB203580 (20 µM, Cell Signaling Technologies 5633), doxorubicin (1 µM, Sigma D1515), and nutlin3a (5 µM, Sigma SML0580),

### Plasmids and cloning

The splicing reporter RG6 construct (Addgene 80167) was amplified (primers 1 and 2) and cloned into the backbone of the pLVX-Bsd-CMV-MDM4 plasmid (Addgene 202669), excluding the *MDM4* insert, using Gibson Assembly [74] generated in-house using primers 3 and 4 to generate a linearized vector.

The *MDM4-S* sequence was generated from the pLVX-Bsd-CMV-MDM4 plasmid using primers 5 and 6. It was cloned into the backbone of the pLVX-Bsd-CMV-MDM4 plasmid using Gibson Assembly with primers 4 and 7 to generate a linearized vector. The *MDM4-S* sequence corresponds to Ensembl transcript ENST00000391947.

All the linearized and amplified products were DpnI-digested before assembling. The assembled products were transformed into DH5α competent cells generated in-house, derived from Library Efficiency DH5α competent cells (Thermo Fisher Scientific 18263012). Correct insertions into the target vector were confirmed by either whole plasmid or Sanger sequencing (Microsynth).

**Table 1:**
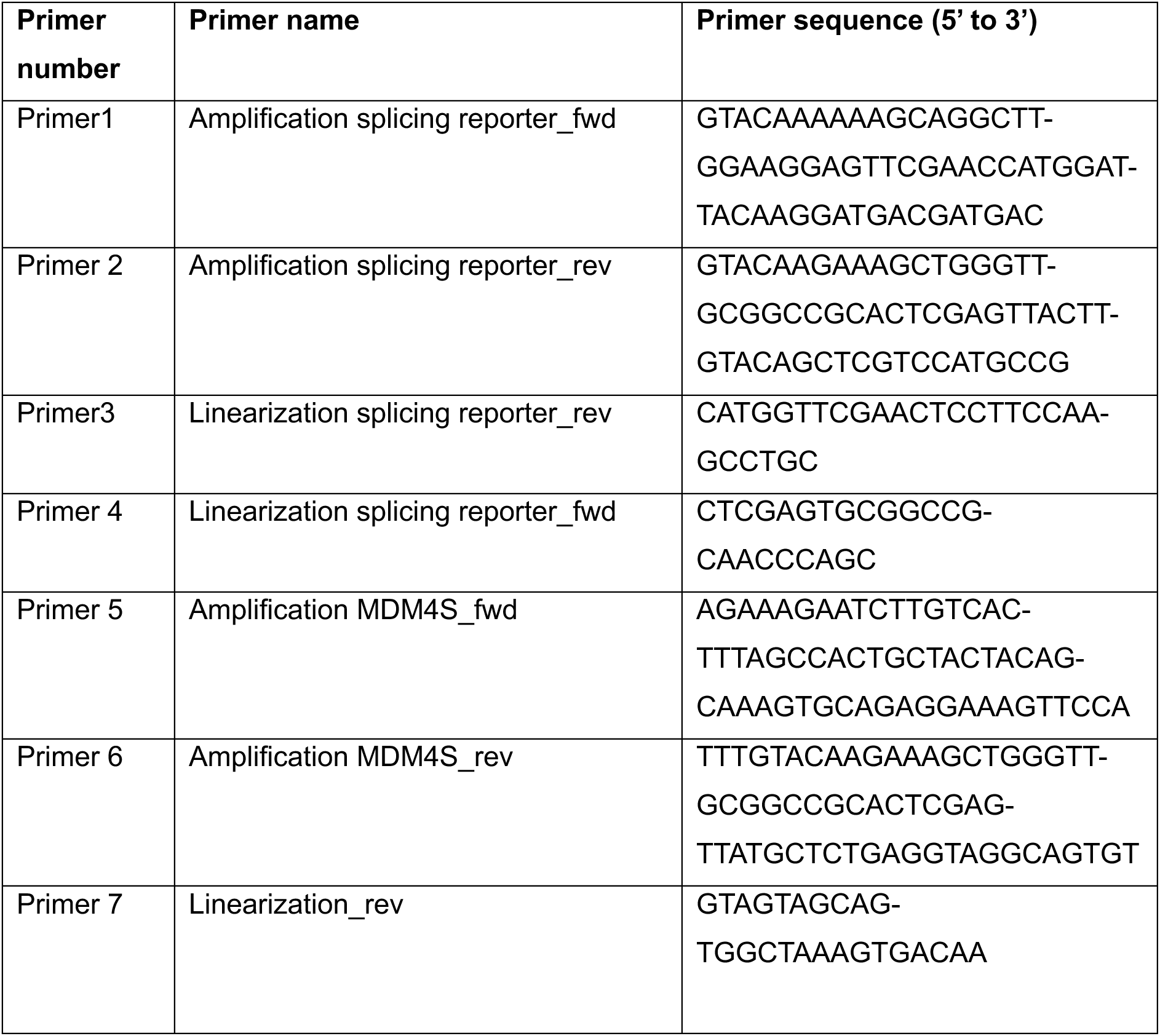
Primers used for Gibson Assembly Cloning in this study.

### siRNA transfections

Cells were reverse-transfected using Lipofectamine RNAiMAX (Invitrogen 13778100) according to the manufacturer’s instructions with siRNAs at a final concentration of 10-20 nM for 24 hours. Knockdown efficiencies were confirmed by western blotting. For G_1_ arrest experiments, transfections were carried out in the constant presence of palbociclib.

**Table 2:** siRNA sequences used in this study.

| siRNA target | Product |
| --- | --- |
| siControl (siGENOME non-targeting siRNA #5) | Dharmacon D-001210-05-05 |
| siMDM4 | Dharmacon L-006536-02-0005 |
| siP21 | Dharmacon J-003471-09-0002 |
| siSRSF2 | Ambion s12730 |

### Lentivirus and stable cell line generation

Lentiviruses were generated as described previously [75]. Briefly, HEK293T cells were co-transfected with a plasmid containing the gene of interest and lentiviral packaging plasmids (pMD2 and psPAX2, gifts from Randall W. King) using Lipofectamine 3000 (Invitrogen L3000015) according to the manufacturer’s protocol. Media was refreshed after 24 hours, and lentiviral supernatant was collected and filtered after an additional 24 hours. RPE1 WT cells were infected with lentiviral supernatants in the presence of 5-10 μg/mL protamine sulfate for 24 hours. Media was refreshed for an additional 24 hours before selecting cells in the appropriate antibiotic-containing media and/or by FACS-sorting cells based on the relevant fluorescent marker(s).

RPE1 p53 knockout cells were generated by directly transfecting RPE1 WT cells with a TP53 sgRNA (CRISPR18517_SGM, Invitrogen A35533) with Cas9 nuclease using Lipofectamine CRISPRMAX (Invitrogen CMAX00001) according to the manufacturer’s instructions. Cells were transfected for 48 hours and were then expanded for an additional 24 hours in fresh media. Following expansion, cells were selected with nutlin3a. Loss of p53 in the resulting polyclonal cell line was confirmed by western blot.

RPE1 RG6 splicing reporter cell line was generated using lentiviral transduction as described above. Cells were selected with 6 µg/mL blasticidin (InvivoGen, ant-bl) for seven days. Cells were subsequently sorted for GFP/dsRED positive cells using a FACS Aria III 5L controlled by the FACS Diva Software and equipped with 488 nm, 561 nm, 633 nm, and 405 nm laser lines using a 100 µm nozzle.

RPE1 MDM4 cDNA cells were generated using lentiviral transduction as described above, with the MDM4 cDNA delivered from pLVX-Bsd-CMV-MDM4. Cells were selected in 6 μg/mL blasticidin for seven days. MDM4 cDNA expression was confirmed by RT-PCR.

### Cell size measurements

Cell sizes were measured using a Multisizer 4e (Beckman Coulter) using Isotone II (Beckman Coulter NC2470899) as a diluent and a 100 μm aperture tube. Trypsinized cells were re-suspended in DMEM/F12 and diluted in 10 mL Isotone II before measuring. Cell volume distributions were calculated using a custom R script.

### RNA sequencing sample preparation

Enlarged and size-constrained RPE1 cells were collected as in **Figure 1A** in quadruplicate. Cell pellets were flash frozen in liquid nitrogen, and RNA was extracted using an RNeasy Mini Kit (Qiagen 74104). Total RNA concentrations were measured using a Qubit Fluorometer (Invitrogen Q33238) and normalized. Set #1 from ERCC ExFold RNA Spike-In Mixes (Invitrogen 4456739) was added to each sample at a 1:1000 mass ratio.

### RNA sequencing analysis

#### Library preparation

Extracted RNA was prepared for sequencing by the Functional Genomics Center Zurich (FGCZ). The quality of the isolated RNA was determined with a Fragment Analyzer (Agilent, Santa Clara, California, USA). The TruSeq Stranded Total RNA Library Prep Gold (Illumina, Inc, California, USA) was used in the successive steps. Briefly, total RNA samples (100-1000 ng) were depleted of ribosomal RNA and then reverse-transcribed into double-stranded cDNA. The cDNA samples were fragmented, end-repaired, and adenylated before ligation of TruSeq adapters containing unique dual indices (UDI) for multiplexing. Fragments containing TruSeq adapters on both ends were selectively enriched with PCR. The quality and quantity of the enriched libraries were validated using a Fragment Analyzer (Agilent, Santa Clara, California, USA). The product was a smear with an average fragment size of approximately 260 bp. The libraries were normalized to 10nM in 10 mM Tris HCl, pH 8.5 with 0.1% Tween 20.

#### Cluster Generation and Sequencing

The NovaSeq X Plus (Illumina, Inc, California, USA) was used for cluster generation and sequencing according to a standard protocol. Sequencing was performed using paired-end reads of 2 x 150bp.

#### Transcript alignment and quantification

Raw reads were first cleaned by removing adapter sequences and poly-x sequences (> 9 nt used for detection) using fastp (Version 0.23.4) [76]. Sequence pseudo alignment of the resulting high-quality reads to the Human reference genome (build GRCh38.p13) and quantification of transcript-level expression (gene model definition from GENCODE release 42) were carried out using Kallisto (Version 0.46.1) [77].

#### Transcript isoform analysis

Differential isoform usage and isoform switching events were analyzed using the IsoformSwitchAnalyzeR package (v2.6.0) in R v4.4.2 [78]. Estimated transcript counts and abundances from Kallisto were imported together with the GENCODE release 42 isoform annotation (exon structure and annotated coding sequences) and the corresponding transcript nucleotide sequences; the counts were used for statistical testing and the abundances for effect-size estimation.

Transcript quantification data were filtered to remove lowly expressed genes (gene expression cutoff = 1 TPM), single-isoform genes, and isoforms contributing less than 1% of their gene’s expression (isoform-fraction cutoff = 0.01). Differential isoform usage was tested using the DEXSeq-based statistical framework implemented in IsoformSwitchAnalyzeR, applied to all isoforms so that isoform-fraction differences (dIF) and FDR-corrected p-values (q-values) were obtained for every isoform independently of any effect-size cutoff. Isoform switches were then called at a significance threshold of alpha = 0.05. To assess how sensitive the called switches are to the effect-size threshold, three differential isoform-fraction cutoffs were compared: dIF = 0.01, 0.05 and 0.10. At each cutoff, the alternative-splicing analysis was performed for the switching genes. A dIF cutoff of 0.05 was used for all analyses shown in this manuscript.

### Tyramide signal amplification (TSA) sequencing

#### TSA labeling reaction

TSA-seq was performed according to the TSA-seq 2.0 protocol, as described previously [34], with modifications for T-300 cell culture flasks. Enlarged cells were seeded at 5.33 x 10^3^ cells/cm^2^ and size-constrained cells were seeded at 3.2 x 10^4^ cells/cm^2^, and 20 million cells were collected per condition. Cells were fixed in 1.6% paraformaldehyde (PFA) in PBS for 20 minutes at room temperature, washed once with PBS, and permeabilized with 0.5% Triton X-100 in PBS (0.5% PBS-T) in three 5-minute incubations. Residual aldehyde groups were quenched with 20 mM glycine using three 5-minute washes in PBS. Endogenous peroxidase activity was quenched with 1.5% H₂O₂ (v/v) in PBS for 1 hour followed by three washes in PBS. Cells were blocked in 5% goat serum (Gibco 16210072) in PBS for 1 hour and incubated with anti-SON primary antibody for 16 hours. Cells were then washed three times with 0.1% PBS-T and incubated with HRP-conjugated secondary antibody for 10 hours. Following additional three washes with 0.1% PBS-T, tyramide-biotin labeling (Sigma-Aldrich, SML2135) was performed for 30 minutes at room temperature under condition E (tyramide-biotin 1:300, 0.0015% H_2_O_2_, 50% sucrose) as previously described [34].

#### Checking labeling efficiency by immunofluorescence

After tyramide labeling, a small fraction of cells was scraped from the flasks and transferred on glass coverslips (Epredia CB00120RA120MNZ0). Cells were incubated with Streptavidin-Alexa Fluor 594 (Biolegend 405240; 1:200) to detect biotinylation and an Alexa Fluor antimouse 488-conjugated secondary antibody to detect SON for 2 hours at room temperature. Cells were washed with 0.1% PBS-T for 5 minutes three times, incubated with 0.1 μg/mL Hoechst 33342 to stain DNA, and mounted using Prolong Diamond Antifade mounting medium (Thermo P36961) before imaging.

#### Cell lysis and proteinase K digestion

Cell were lysed in high TE buffer supplemented with 1% SDS and 0.2 mg/mL proteinase K (Thermo Scientific, EO0491). Lysates were transferred to conical tubes, supplemented with 0.05 mg/mL glycogen (Roche, 10901393001), and incubated at 55°C for 16 hours in a hybridization chamber with constant rotation. Samples were stored at −80°C.

#### Genomic DNA extraction

Samples were reverse crosslinked by incubating at 65°C for 24 hours with constant rotation after adding NaCl to the lysate at a final concentration of 0.2 M.

Genomic DNA was extracted by adding an equal volume of Phenol:Chloroform:Isoamyl alcohol (25:24:1, Fisher BioReagents BP1752I-44), vortexing for 20 seconds, and centrifuging for 5 minutes at maximum speed. The aqueous layer was collected and the extraction was repeated once. Next, an equal volume of Chloroform:Isoamyl alcohol (24:1, Sigma C0549) was added, vortexed vigorously for 20 seconds, and centrifuged at maximum speed for 5 minutes. The aqueous layer was collected in a new tube and incubated open at 55 °C for 1 hour to evaporate the phenol. After cooling, RNase A (New England Biolabs T3018L) was added at a final concentration of 25 µg/mL and rotated overnight at 37°C. The genomic DNA isolation process was repeated once more before DNA precipitation.

For DNA precipitation, 1/10 volume of 3M Sodium acetate (pH 5.2) and glycogen (Roche 10901393001) to a final concentration of 0.05 mg/mL was added to the sample. Next, two volumes of 100% EtOH were added and incubated at −20°C for 2 hours. Samples were centrifuged in conical tubes at maximum speed at 4°C for 15 minutes, and the supernatant was discarded. The pellet was rinsed with ice-cold 70% EtOH and centrifuged for 5 minutes at maximum speed at 4°C. The supernatant was discarded, and the pellet was air dried before dissolving in nuclease-free water.

#### Genomic DNA fragmentation

Genomic DNA was sonicated using a Bioruptor Pico (Diagenode) with 30 seconds ON and 30 seconds OFF pulses. Fragmentation was monitored by agarose gel electrophoresis and performed until a DNA smear in the 100–600 bp range was observed.

#### Checking biotinylation by dot blot

Dot blot was performed using standards described in [34]. The standards and the samples (1.5 µL) were blotted onto a nitrocellulose membrane (0.45 µm, Biorad 1620115), dried, and UV crosslinked at 0.24J using UV Stratalinker 2400 (Agilent Technologies). The membrane was blocked with Superblock blocking buffer (Thermo 37515) with 0.05% Tween-20 at room temperature for 1 hour. The membrane was then incubated overnight with streptavidin -HRP (Invitrogen 43-4323) at 1:10,000 in blocking buffer at 4°C. The membrane was washed six times with 0.05%Tween-20 in TBS at room temperature for 5 minutes before developing with SuperSignal West Femto chemiluminescent substrate (Thermo 34094) on an iBright imaging system (Invitrogen).

#### Isolation, extraction, and precipitation of biotinylated DNA

A spike-in from Drosophila DNA containing BAC was made as described in [34]. 0.2 ng of biotinylated and non-biotinylated DNA were added to all samples. A 10 µL aliquot of the sample was used for input library preparation.

An equal volume of 2x wash and binding buffer (10 mM Tris-Cl pH 7.5, 1 mM EDTA, 2M NaCl) was added to the samples. M270 Streptavidin Dynabeads (Invitrogen 65306) were incubated with DNA for 30 minutes at room temperature and 2 hours at 4°C with constant rotation. Beads were washed four times with 1x wash and binding buffer, four times with TSE500 buffer (20 mM Tris pH 8.0, 1% Triton X100, 0.1% SDS, 2 mM EDTA, 500 mM NaCl), four times again with 1x wash and binding buffer and, finally, four times with high TE buffer. Beads were then incubated in high TE buffer at a 6:1 ratio with 0.5% SDS, 1 mg/mL proteinase K (New England Biolabs P8107S) and free biotin (30 µL of 0.5 mg/mL per 100 µL of beads) at 55°C for 16 hours with constant rotation to release biotinylated DNA from the beads.

The bead-digested solution was brought to a final concentration of 0.2M NaCl. Equal volumes of Phenol:Chloroform:Isoamyl alcohol (25:24:1) were added, and samples were vortexed for 20 seconds and spun down for 5 minutes at maximum speed. The aqueous layer was collected, and the process was repeated. Next, an equal volume of Chloroform:Isoamyl alcohol (24:1, Sigma C0549) was added to each sample, vortexed for 20 seconds and centrifuged at maximum speed for 5 minutes. The aqueous layer was collected and incubated at 55°C for 1 hour to allow residual phenol to evaporate. Biotinylated DNA was precipitated by adding 1:10 volume of 3M Sodium citrate (pH 5.2) and glycogen (Roche 10901393001) to a final concentration of 0.05 mg/mL. Next, two volumes of 100% ethanol were added to the samples and incubated at −20°C for 2 hours. Samples were centrifuged at maximum speed at 4°C for 15 minutes and the supernatant was discarded. Pellets were rinsed with ice-cold 70% ethanol and centrifuged for 5 minutes at maximum speed at 4°C. The supernatant was discarded, and the pellet was air dried before dissolving in Elution Buffer (EB) (Qiagen 19086). DNA concentrations were determined by Qubit (Invitrogen Q32854).

#### Library preparation

Libraries were made with 2.75 - 7.5 ng of DNA (biotinylated DNA for pulldown and corresponding genomic DNA kept aside before pulldown for input) using the NEBNext® Multiplex Oligos for Illumina® (Index Primers Set 1, E7335S) and NEBNext® UltraTM II DNA Library Prep Kit for Illumina® (E7645S) following manufacturer’s instructions. The library concentration was estimated by qPCR and pooled and sequenced in one lane by single-end sequencing for 100bp read length in a NovaSeq X 10B flow cell.

### TSA-seq data analysis

The FASTQ files generated were aligned to the hg38F reference genome assembly using bowtie2 [79]. The rmdup command in samtools was used to remove PCR duplicates. We used bedtools to divide the genome into 20kb bins and count the reads mapped to each bin for both pulldown and input samples. Bins with fewer than 4 reads in the input were removed.

TSA-seq scores for each bin were calculated as follows: N_pulldown_ is the number of reads mapped to a particular bin in pulldown sample; N_input_ is the number of reads mapped to a particular bin in input samples; N_pulldown_mean_ is the mean of the number of reads mapped to all retained bins in the pulldown; N_input_mean_ is the mean of the number of reads mapped to all retained bins in input. Calculating *N_TSA_’* = (*N_pulldo_*_w*n*_/*N_pulldo_*_w*n*_*mean*_) ⁄ (*N_input_*/*N_input_*__*mean*_), the TSA-seq score = log_2_ *N_TSA_’*.

TSA-seq scores were smoothed using a Hanning window as previously described [34], accessible at https://github.com/lgchang27/TSA-Seq-2020v2. The TSA-seq score for all genes in GENCODE release 47 was extracted from the normalized scores by taking the weighted mean of the TSA-seq scores of the bins which the gene occupies with respect to the length of the bin occupied by the gene.

### Replication timing analysis

Replication timing data for hTERT-RPE1 cells (two-fraction early/late Repli-seq, two replicates per fraction) were obtained from the 4D Nucleome Data Portal (experiment set 4DNESLKJF5N2; generated by the laboratory of David M. Gilbert) [80, 81]. RPKM-normalized read counts in 5-kb bins (4DNFI9R1MQ7U, 4DNFIRNKABQJ, 4DNFIBKO9LI7, and 4DNFIK2TA4DS) were used without further read-level processing. Early and late counts were summed across replicates and quantile-normalized against each other. Replication timing was calculated per gene as log₂(early/late) over ±50 kb around the transcription start site and expressed as a percentile across all genes.

### Immunofluorescence staining

Cells were plated on glass coverslips (Epredia CB00120RA120MNZ0) or in a black-walled 96-well imaging plate (Ibidi 89626) and allowed to attach for 24 hours before staining. Coverslips or plate wells were washed once briefly in PBS and then fixed in 3% paraformaldehyde (PFA) for 15 minutes. Fixed cells were then permeabilized with 0.25% Triton X-100 in PBS for 10 minutes and washed three times for 5 minutes with 0.05% Triton X-100 in PBS. Cells were then blocked in 1% BSA in PBS for 30 minutes and incubated with primary antibodies diluted in 1% BSA in PBS for 1 hour. Cells were washed three times for 5 minutes with 0.05% Triton X-100 in PBS before incubation with the appropriate Alexa Fluor-conjugated secondary antibodies in 1% BSA in PBS for 1 hour. Stained cells were then washed once in 0.05% Triton X-100 in PBS for 5 minutes, then twice in 0.05% Triton X-100 + Hoechst 33342 (0.1 µg/mL) in PBS for 5 minutes. Coverslips were washed briefly in MilliQ water before mounting on glass slides using either Vectashield mounting medium (Vector Laboratories H-1200-10) or ProLong Diamond Antifade mountant (Invitrogen P36961), and 96-well plates were stored and imaged in PBS. All steps were carried out at room temperature.

**Table 3:** Antibodies used for immunofluorescence and TSA-seq in this study.

| Antibody | Manufacturer |
| --- | --- |
| p21 (rabbit, 1:500) | Abcam, ab109520 |
| p53 (mouse, 1:500) | Invitrogen, AHO0152 |
| SC35 (mouse, 1:400) | Merck (Sigma), S4045 |
| SON (mouse, 1:1000) | Santa Cruz Biotechnology, SC-398508 |
| Smith antigen antibody (mouse, 1:200) | Sigma Aldrich, MABF2793 |
| Alexa Fluor 488 goat anti-mouse (mouse, 1:1000) | Thermo Fisher Scientific, A11029 |
| Horse anti-mouse IgG antibody (H+L), peroxidase (0.4 µg/mL) | Vector Laboratories, PI-2000 |
| Alexa Fluor 488 goat anti-rabbit (1:1000) | Thermo Fisher Scientific, A11034 |
| Alexa Fluor 647 goat anti-mouse (1:1000) | Thermo Fisher Scientific, A21235 |
| Alexa Fluor 647 goat anti-rabbit (1:1000) | Thermo Fisher Scientific, A21244 |
| Alexa Fluor 568 goat anti-mouse (1:1000) | Thermo Fisher Scientific, A11031 |
| Alexa Fluor 568 goat anti-rabbit (1:1000) | Thermo Fisher Scientific, A11036 |

### EdU staining

To assess the fraction of G_1_-released cells that enter S-phase over a given time frame, cells were plated on coverslips under G_1_-arresting conditions for 24 hours. Cells were then washed three times to remove drugs from the media and released into fresh media containing 20 µM EdU-alkyne for the indicated amount of time. Detection of EdU-labeled DNA was carried out using Click-iT chemistry for 30 minutes in the dark following a standard protocol (Click-iT EdU Alexa Fluor Imaging Kit, Thermo C10337). DNA was subsequently stained with Hoechst 33342 (0.1 µg/mL) and mounted on glass slides as previously indicated.

### Fixed cell imaging

Fixed cell microscopy was performed on a Nikon Ti2 Eclipse inverted microscope equipped with a 100x/NA 1.45 Plan Apo oil objective, a 40x/NA 0.95 Plan Apo air objective, or a 20x/NA 0.75 PlanApo air objective depending on the experiment. Fluorescence was excited with a Lumencor SpectraX light engine. Hoechst fluorescence was imaged using a 395/25 nm excitation filter and a 432/36 nm emission filter; AlexaFluor488 fluorescence was imaged using a 470/24 nm excitation filter and a 515/30 nm emission filter; AlexaFluor568 fluorescence was imaged using a 575/25 nm excitation filter and a 595/31 nm emission filter; Cy5 fluorescence was imaged using a 640/18 nm excitation filter and a 698/70 nm emission filter. Images were acquired with a Hamamatsu Orca Fusion CoaXpress sCMOS camera controlled by NIS (Nikon) Elements image acquisition software. Five to ten fields of view were collected per condition.

### Live cell video microscopy of FUCCI reporters

G_1_ arrested cells were plated in the presence of palbociclib on an 8-well continuous coverslip dish (Ibidi 80826) for 24 hours to attach. To measure cell cycle release dynamics, cells were washed three times and imaged immediately following the addition of fresh media. Dishes were mounted on a motorized stage (Prior ProScan HLD117NN) housed inside a covered cage microscope incubator (Okolabs) with temperature (37°C) and humidity control in the presence of 5% CO_2_. All images were collected using a Nikon Ti2 motorized inverted microscope equipped with a piezo z-drive (Prior), a 20x/0.75 NA S Fluor air objective, and the Perfect Focus system (Nikon). mCherry fluorescence was excited with a Lumencor Spectra III light engine using a 578/21 nm excitation filter and 641/75 nm emission filter. mAG1 fluorescence was excited using a 474/27 nm excitation filter and a 515/30 nm emission filter. Images were acquired with a Hamamatsu ORCA Fusion BT sCMOS camera controlled by NIS (Nikon) Elements image acquisition software. Brightfield and fluorescence images were captured at 5- to 6-minute intervals; at least four fields of view were collected per condition.

### Image analysis

For fixed cell imaging, all quantifications were performed using standard tools in FIJI [82]. Nuclei were segmented using the DAPI channel, and mean intensity measurements for the indicated antibodies were made as a proxy for nuclear protein concentration since this accounts for changes in nuclear volume. Poorly segmented cells and cells that were at the border of the image were excluded from downstream analyses. Foci were identified using the “Find Maxima” tool and were automatically counted per segmented nucleus.

For live cell imaging of FUCCI markers, cells were randomly selected on the first frame of the video and tracked manually using the same display settings for all videos in the same experiment to insure consistent cell cycle phase calling. Cells that exited the field of view during the duration of imaging were excluded from downstream analyses. Note that, because FUCCI markers are a proxy for the activity of cell cycle-regulated ubiquitin ligases [19], the actual timing of each cell cycle phase may vary, which is why S-phase and G_2_ timing were measured together.

### RT-PCR and qRT-PCR

RNA extraction was performed using the RNeasy Mini Kit (Qiagen 74104) with the RNAse-free DNase set (Qiagen 79254) according to the manufacturer’s instructions. RNA was primed with random hexamers (Roche 11034731001) and reverse-transcribed into cDNA using MultiScribe Reverse Transcriptase (Applied Biosystems 4308228).

Quantitative real-time PCR (qRT-PCR) was performed using the KAPA SYBR FAST kit suitable for qPCR (KAPA Biosystems KK4618) on a Rotor-Gene Q system (Qiagen). All samples were analyzed in technical triplicate for each biological replicate and normalized to ACTB levels.

**Table 4:**
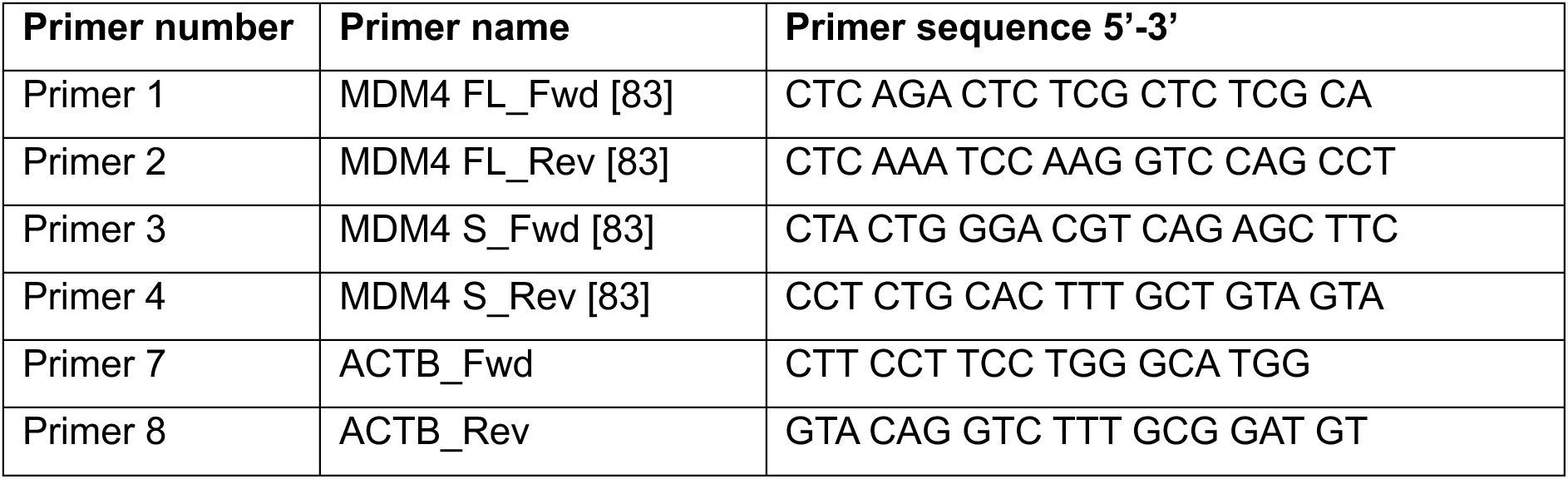
qRT-PCR primers used in this study.

**Table 5:**
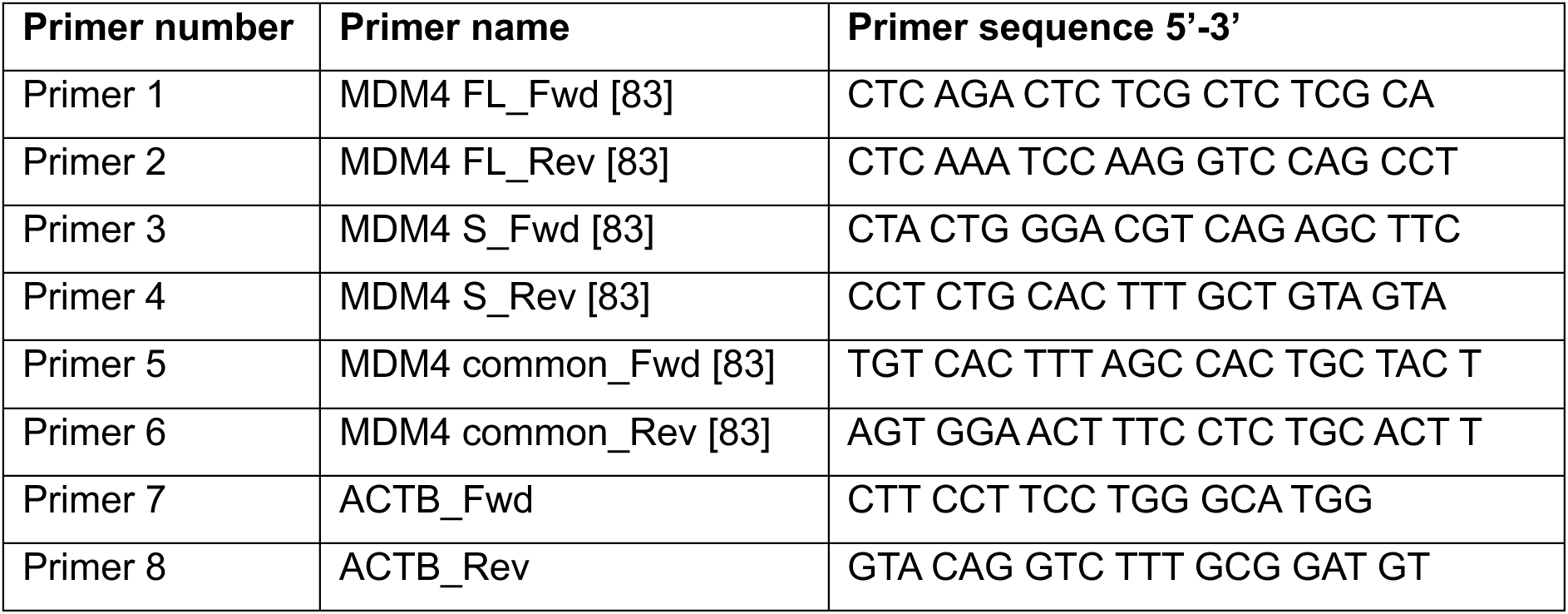
RT-PCR primers used in this study.

### Western blotting

Cells were lysed in RIPA lysis buffer (Invitrogen 89900 or homemade) supplemented with 1x protease/phosphatase inhibitor tablets (Thermo A32961) by periodic vortexing on ice. Lysates were clarified by centrifugation at 16,000 x *g* for 10 minutes at 4°C. Lysates were transferred to fresh tubes, and protein concentrations were measured using BCA assay (Pierce 23225) according to the manufacturer’s instructions. Lysates were mixed with 4x LDS sample buffer (Invitrogen NP0007) to a final concentration of 1x and were heated to 95°C for 10 minutes. Equal masses of protein were loaded on Bolt or NuPAGE 4-12% Bis-Tris gels (Invitrogen) and resolved in either 1x MOPS or 1x MES SDS running buffer (Invitrogen B000102 or B0002). Proteins were transferred to PVDF membrane at 250-350 mA at 4°C for 1 hour and 15 minutes using a wet transfer apparatus (BioRad 1703930). Membranes were blocked in 5% non-fat dry milk in TBS-T before incubating with primary antibodies in 5% milk in TBS-T overnight with agitation. Membranes were washed three times in TBS-T for ∼10 minutes each, incubated with a secondary mouse or rabbit IgG HRP conjugated antibody (Vector Laboratories PI-2000 or PI-1000-1, 1:8000) in 5% milk for 45-60 minutes, washed again, and then developed using enhanced chemiluminescent substrate solutions as described by the manufacturer (Thermo 34095, 344578) on a Fusion imaging system (Vilber).

**Table 6:** Antibodies used for western blot.

| Antibody | Manufacturer |
| --- | --- |
| p21 (rabbit, 1:1000) | Cell Signaling Technology, 2947 |
| GAPDH (mouse, 1:3000) | Abcam, ab8245 |
| PTBP1 (rabbit, 1:1000) | Cell Signaling Technology, 57246 |
| U2AF2 (rabbit, 1:1000) | Invitrogen, PA5-30442 |
| CCAR1 (rabbit, 1:1000) | Abcam, ab70243 |
| hnRNPK (rabbit, 1:1000) | Cell Signaling, 9081 |
| p53 (mouse, 1:500) | Santa Cruz, sc-126 |
| MDM4 (rabbit, 1:1000) | Bethyl, A300-287A |

### Colony formation assay

Cells were seeded at ∼200 cells/cm^2^ in the presence of palbociclib for 24 hours. Cells were then washed 3x in media and released into fresh media for 10 days. Cells were then washed with ice-cold PBS followed by incubation with ice-cold 100% methanol for 10 minutes on ice. Cells were then stained with 0.5% crystal violet (w/v) in 20% ethanol for 10 minutes at room temperature followed by washing in DI water until clear. Plates were dried at room temperature before imaging. Colony formation (percentage of the well surface covered) was quantified using the automated ColonyArea macro in ImageJ [84].

### Gene enrichment analysis

Gene enrichment analysis was carried out on the indicated gene sets using the ShinyGO tool [85] for biological processes as indicated in [9].

### Statistical methods

No statistical methods were used to predetermine sample size. Statistical tests (where relevant) are indicated in the figure legend along with the corresponding p-values and whether multiple comparisons corrections were used. In all other experiments (except where indicated in the *Image Analysis* section), no data were excluded from analyses. Apart from time-lapse video microscopy, no samples were measured repeatedly, and measurements were taken from distinct samples.

## Data Availability

Repli-seq data are available from the 4DN Data Portal (https://data.4dnucleome.org) under accession 4DNESLKJF5N2. Phospho-proteomic data are available in our contemporaneous work [22]. TSA-seq and RNA-seq data have been deposited to GEO and will be made available upon publication.

## Acknowledgements

We are grateful to R.W. King for sharing the hTERT-RPE1 FUCCI cell line. We thank T. Cooper, R.W. King, and J. Sheltzer for plasmids and J. Corn for antibodies used in this study. We thank the laboratory of David M. Gilbert and the 4D Nucleome Network for making the hTERT-RPE1 Repli-seq data available. We thank the Scientific Center for Optical and Electron Microscopy (ScopeM, ETH Zürich) and the Center for Microscopy and Image Analysis (ZMB, University of Zürich) for assistance with microscopy experiments and the Functional Genomics Center Zürich (FGCZ) for support with the RNA sequencing experiment. This work was funded by an SNSF Ambizione grant (PZ00P3_223940), a Novartis Foundation research grant (24C253), and an ETH Career Seed grant (23-2 Seed-013) to S.M.

## Author Contributions

Conceptualization: S.M.; Methodology: K.W., F.N., S.M.; Investigation and Formal Analysis: L.M., K.W., S.N., L.B., F.N., S.M.; Resources: G.E.N., M.A.; Writing – Original Draft: S.M.; Writing – Reviewing & Editing: All authors; Funding Acquisition: S.M.; Supervision: R.S., A.S.B., M.A., S.M.; Project Administration: S.M.

## Supplementary Figures

**Figure S1: Related to Figure 1.**
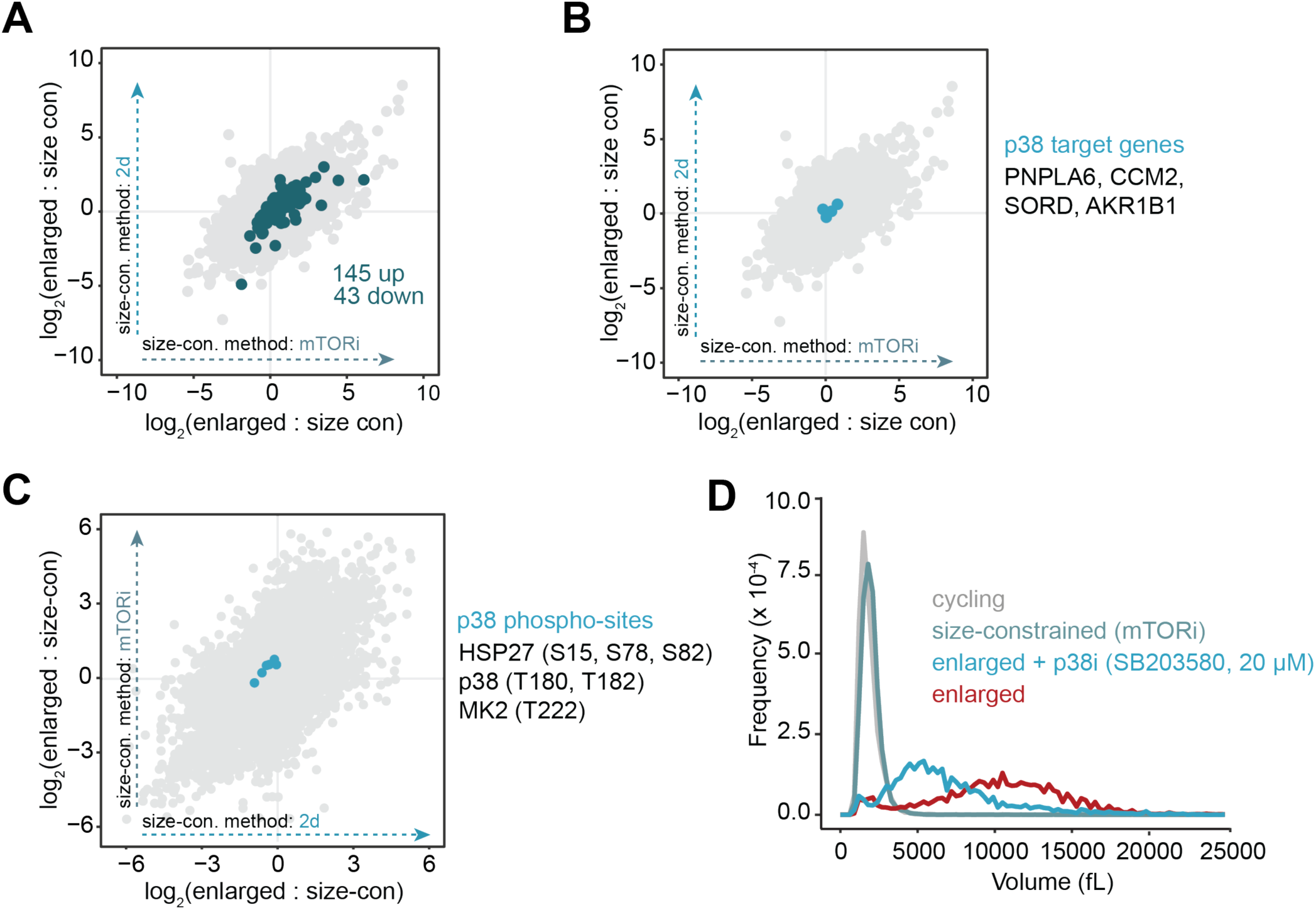
**A.** Comparison of all transcript abundances in enlarged cells compared to each method of size-constraint (2d on the y-axis; mTORi on the x-axis). Annotated p53 target genes that are significantly changing (FDR < 0.01) are shown in blue, with 142 upregulated and 43 downregulated out of an annotated dataset of 342. **B.** The same figure depicted in (**A**) with annotated p38 target genes highlighted instead of p53 target genes. p38 target genes were obtained from Crozier, et al. *Mol Cell* (2023). **C.** Quantification of phospho-peptide abundance in enlarged vs. size-constrained cells (2d on the x-axis; mTORi on the y-axis). The indicated p38 target phospho-peptides are indicated in blue. Data were obtained from our contemporaneous work Wentinck, et al. *bioRxiv* (2026). **D.** Example cell volume measurements for cycling, size-constrained (mTORi), and enlarged cells +/- p38 inhibitor SB203580. Enlarged cells that were treated with SB203580 were continuously treated during the palbociclib-mediated arrest before measuring cell volume.

**Figure S2: Related to Figure 2.**
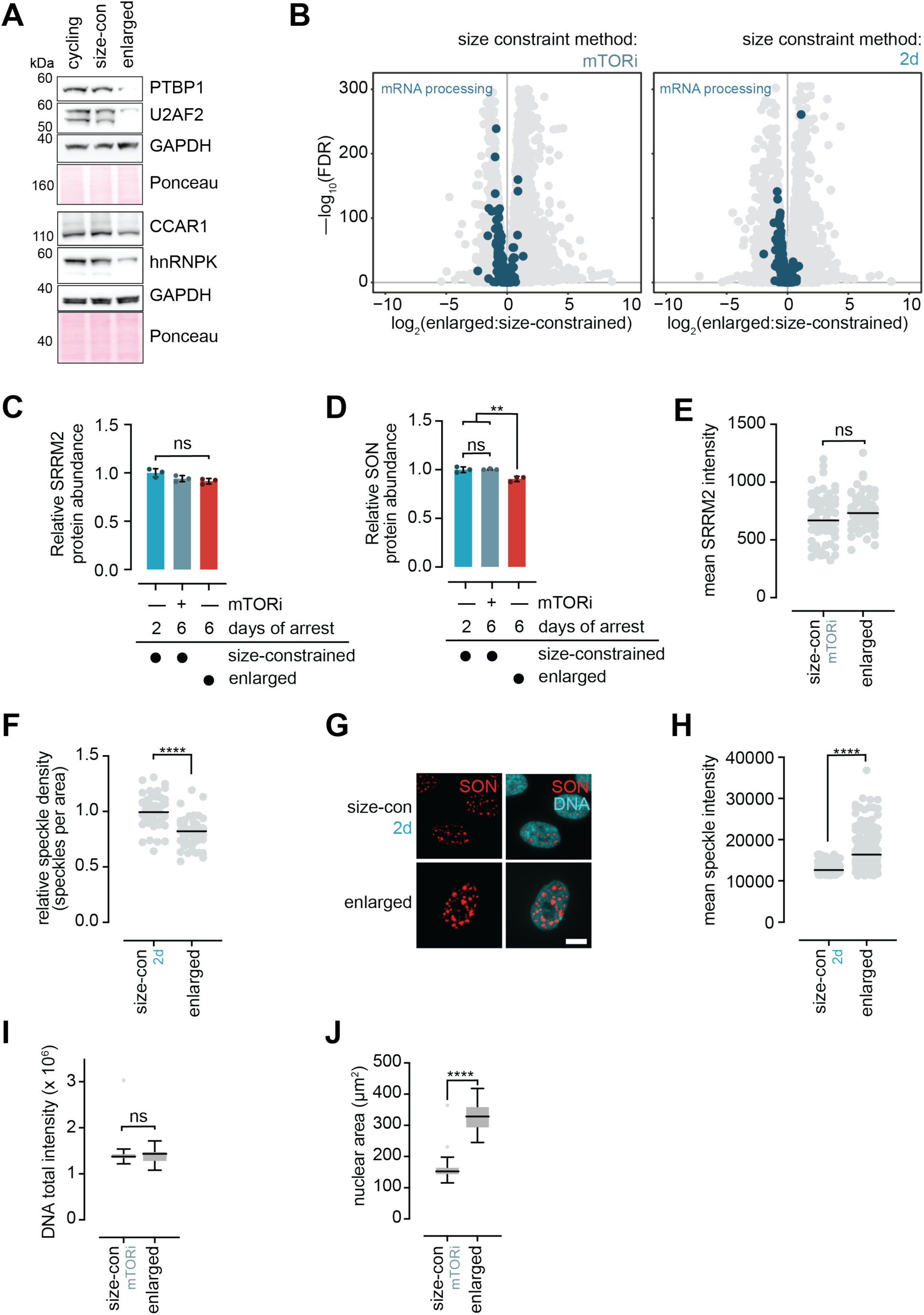
**A.** Western blots of the indicated mRNA processing proteins in cycling, size-constrained (mTORi), and enlarged cells. GAPDH and Ponceau staining are shown as loading controls. **B.** Transcriptomic data for enlarged vs. size-constrained cells (mTORi on left; 2d on right). Genes that fall under the mRNA processing GO category (GO: 0006397) are highlighted in blue. **C.** Relative SRRM2 protein abundance in enlarged and size-constrained cells obtained by proteomics from Manohar, et al. *Mol Cell* (2023). ns : p ≥ 0.06 by one-way ANOVA followed by Tukey’s multiple comparisons test. **D.** Relative SON protein abundance in enlarged and size-constrained cells obtained by proteomics from Manohar, et al. *Mol Cell* (2023). ns : p = 0.97; ** : p ≥ 0.0045 by one-way ANOVA followed by Tukey’s multiple comparisons test. **E.** Quantification of mean SRRM2 intensity per cell in size-constrained (mTORi) and enlarged cells. Bar indicates median value. ns: p = 0.083 by unpaired, two-tailed t-test. n ≥ 56 cells per condition. **F.** Quantification of relative speckle density (speckles per nuclear area) in enlarged vs. size-constrained (2d) cells. Bar indicates median value. **** : p < 0.0001 by unpaired, two-tailed t-test. n ≥ 40 cells per condition. **G.** Example images of the experiment shown in (**F**). Scale bar = 10 µm. **H.** Quantification of mean intensity per speckle in the experiment shown in (**F**) and (**G**). Bar indicates median value. **** : p < 0.0001 by unpaired, two-tailed t-test. n ≥ 40 cells per condition. **I.** Tukey plots quantifying total DNA intensity in enlarged and size-constrained (mTORi) cells. ns : p = 0.57 by unpaired, two-tailed t-test. n = 46 cells per condition. **J.** Tukey plots quantifying nuclear area for the experiment shown in (**I**). **** : p < 0.0001 by unpaired, two-tailed t-test.

**Figure S3:**
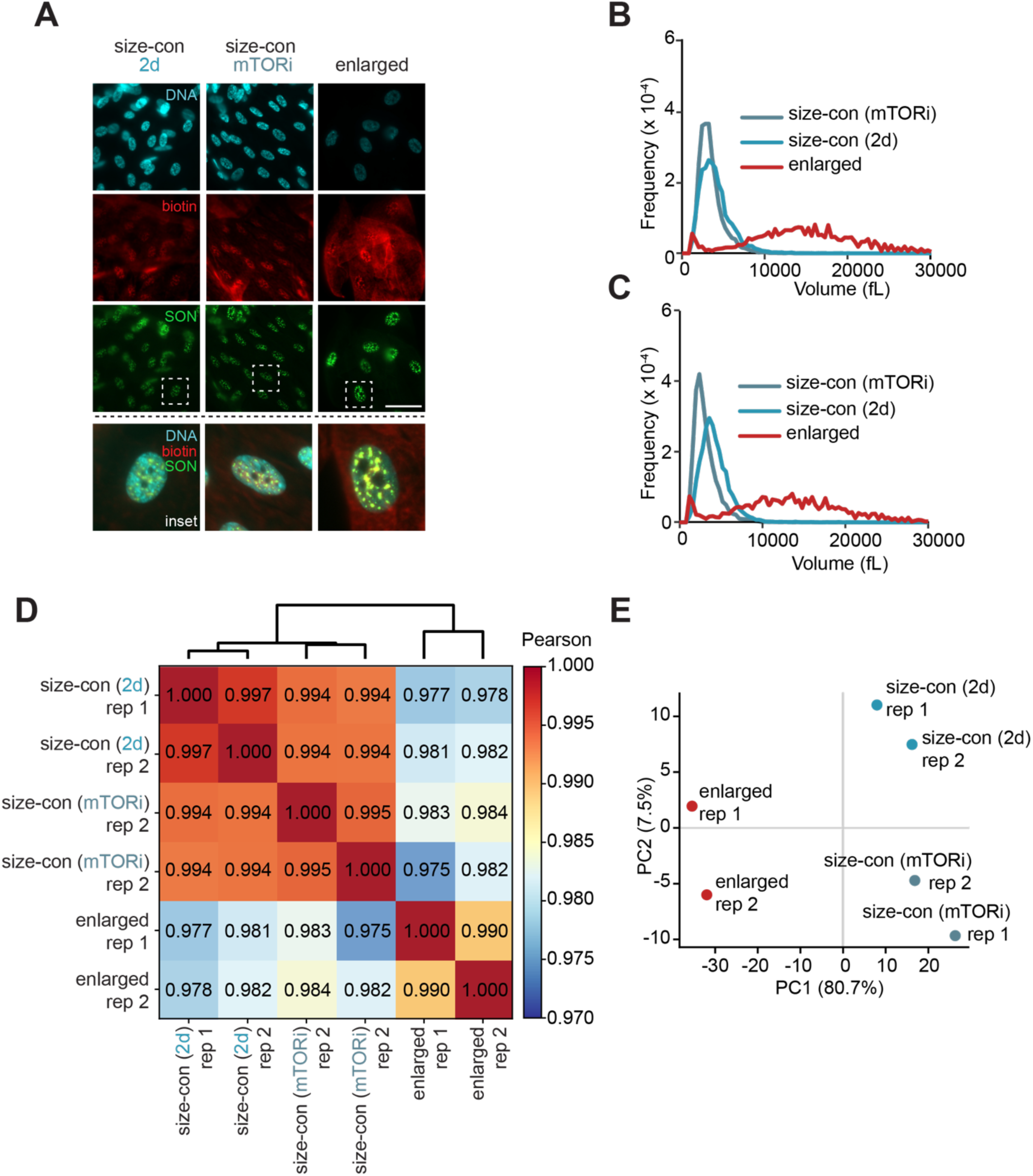
Quality control data for TSA-seq. Related to Figure 3. **A.** Representative images from TSA labeling check in enlarged and size-constrained cells, as detailed in Materials and Methods. Scale bar = 50 µm. **B.** Cell volume measurements for biological replicate 1 for SON TSA-seq. **C.** Cell volume measurements for biological replicate 2 for SON TSA-seq. **D.** Heat map of the Pearson correlation values for each of the conditions and replicates in the SON TSA-seq experiment. **E.** Principal component analysis (PCA) for the conditions and replicates of the SON TSA-seq experiment.

**Figure S4: Related to Figure 3.**
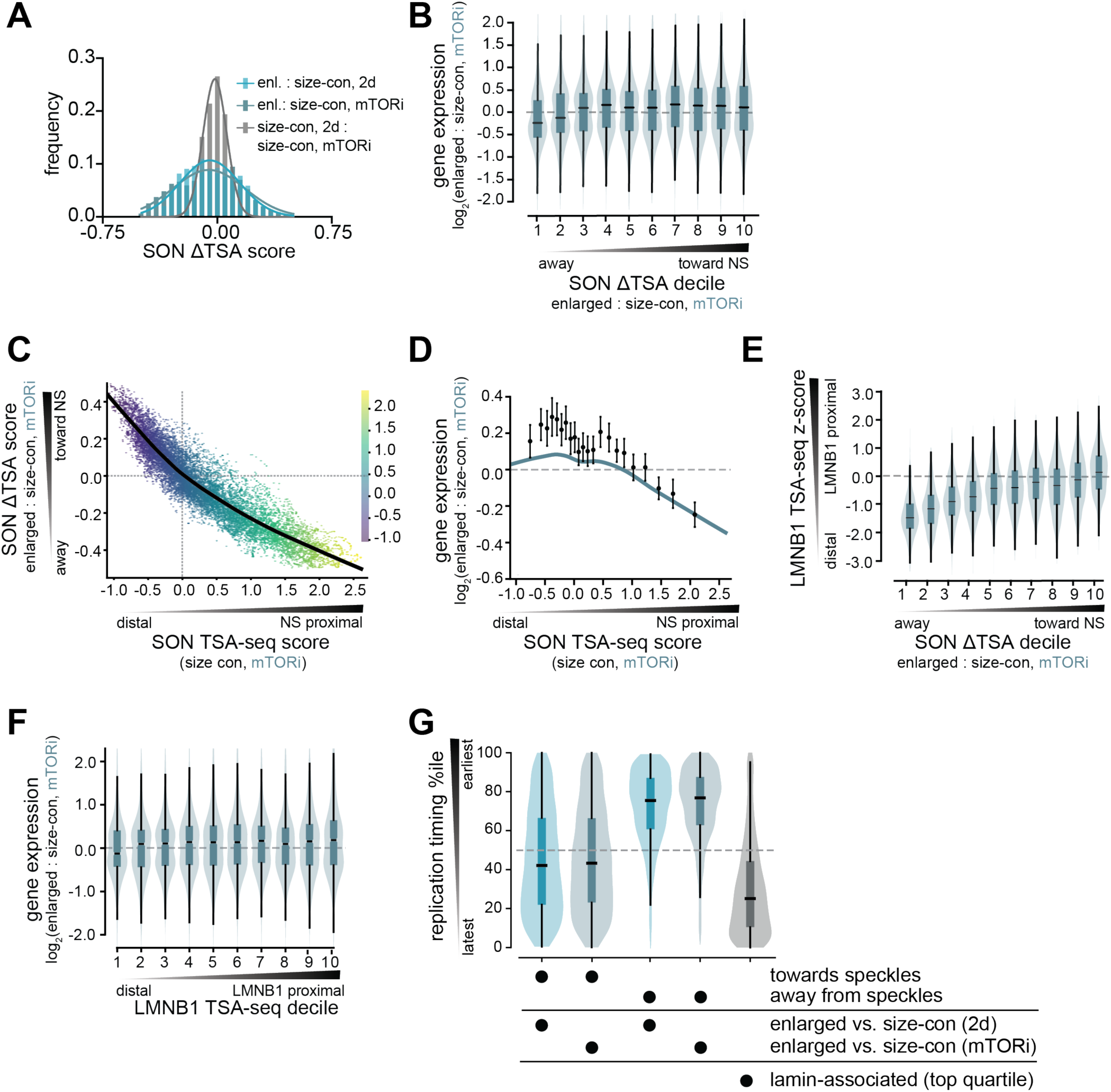
**A.** Distribution of SON ΔTSA values for the indicated comparisons. **B.** Violin plots depicting gene expression for significantly changing (FDR < 0.05) genes in each SON ΔTSA decile for enlarged vs. size-constrained cells (mTORi). Spearman’s ρ = 0.10. n = 10,602 genes. **C.** Scatter plot depicting the change in SON ΔTSA score between enlarged and size-constrained cells (mTORi) as a function of the SON TSA-seq score in size-constrained cells (mTORi) for each gene. Spearman’s ρ = −0.92. Color scale indicates the SON TSA-seq score in size-constrained cells (mTORi). Black curve depicts the LOWESS fit. **D.** Gene expression for significantly changing (FDR < 0.05) genes between enlarged and size-constrained cells (mTORi) plotted as a function of the SON TSA-seq score in size-constrained cells (mTORi). Spearman’s ρ = −0.12. Individual data points depict the mean of 22 equally sized gene bins, and error bars represent 95% confidence intervals. Blue line depicts the LOWESS fit. n = 10,602 genes. **E.** Violin plots depicting the Lamin B1 TSA-seq z-score for each SON ΔTSA decile for enlarged vs. size-constrained cells (mTORi). Spearman’s ρ = 0.47. n = 12,294 genes. **F.** Violin plots depicting gene expression changes in enlarged cells vs. size-constrained cells (mTORi) for significantly changing (FDR < 0.05) genes in each LMNB1 TSA-seq decile. Spearman’s ρ = 0.06. n = 9,099 genes. **G.** Violin plots depicting replication timing percentiles for genes moving towards or away from nuclear speckles (top and bottom quartile of SON ΔTSA among genes with SON TSA-seq score > 0) in enlarged vs. size-constrained cells (2d or mTORi) and for lamin-associated genes (top quartile of LMNB1 TSA-seq z-score). p = 1.4 x 10^-240^ (2d) and 1.1 x 10^-242^ (mTORi) for towards vs. away by two-sided Mann–Whitney U test. Dashed line depicts the median replication timing of all genes.

**Figure S5: Related to Figure 4.**
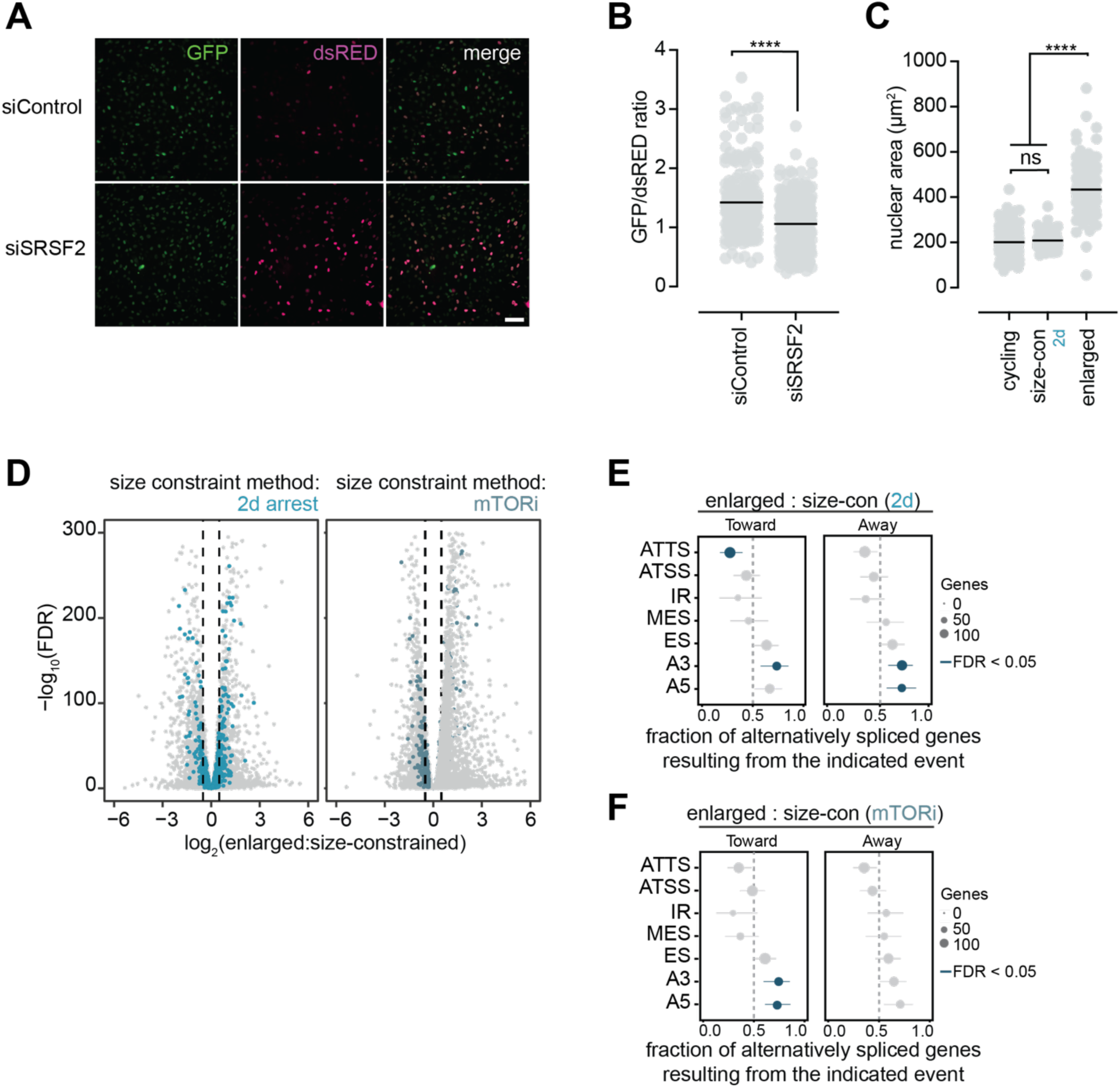
**A.** Example images of the RG6 splicing reporter RPE1 cell line transfected with an siRNA directed against the mRNA splicing regulator SRSF2. Scale bar = 100 µm. **B.** Quantification of the GFP/dsRED ratio per cell for the experiment shown in (**A**). Bar indicates median value. **** : p < 0.0001 by two-tailed, unpaired t-test. n ≥ 280 cells per condition. **C.** Example nuclear area measurements for the experiment shown in **Fig 4B-4C**. Bar indicates median value. **** : p < 0.0001; ns : p = 0.33 by one-way ANOVA followed by Tukey’s multiple comparisons test. n ≥ 170 cells per condition. **D.** Volcano plots indicating total gene expression changes for all isoforms of a given transcript in enlarged vs. size-constrained cells (2d on the left; mTORi on the right). Genes that undergo isoform switching events are highlighted in blue. **E.** Enrichment of alternative splicing events among genes with isoform switches that move towards or away from nuclear speckles (top and bottom quartile of SON ΔTSA scores among genes with SON TSA-seq score > 0, respectively) for enlarged vs. size-constrained cells (2d). Data points depict the fraction of genes in which the isoform gaining usage in enlarged cells carries the indicated event. Error bars indicate 95% confidence interval, and point size indicates number of genes. Blue data points indicate an FDR < 0.05 by two-sided binomial test with Benjamini–Hochberg correction. ATTS: alternative transcription termination site, ATSS: alternative transcription start site, IR: intron retention, MES: multiple exon skipping, ES: exon skipping, A3: alternative 3′ splice site and A5: alternative 5′ splice site. **F.** As shown in (**E**), but comparing enlarged vs. size-constrained cells (mTORi).

**Figure S6: Related to Figure 5.**
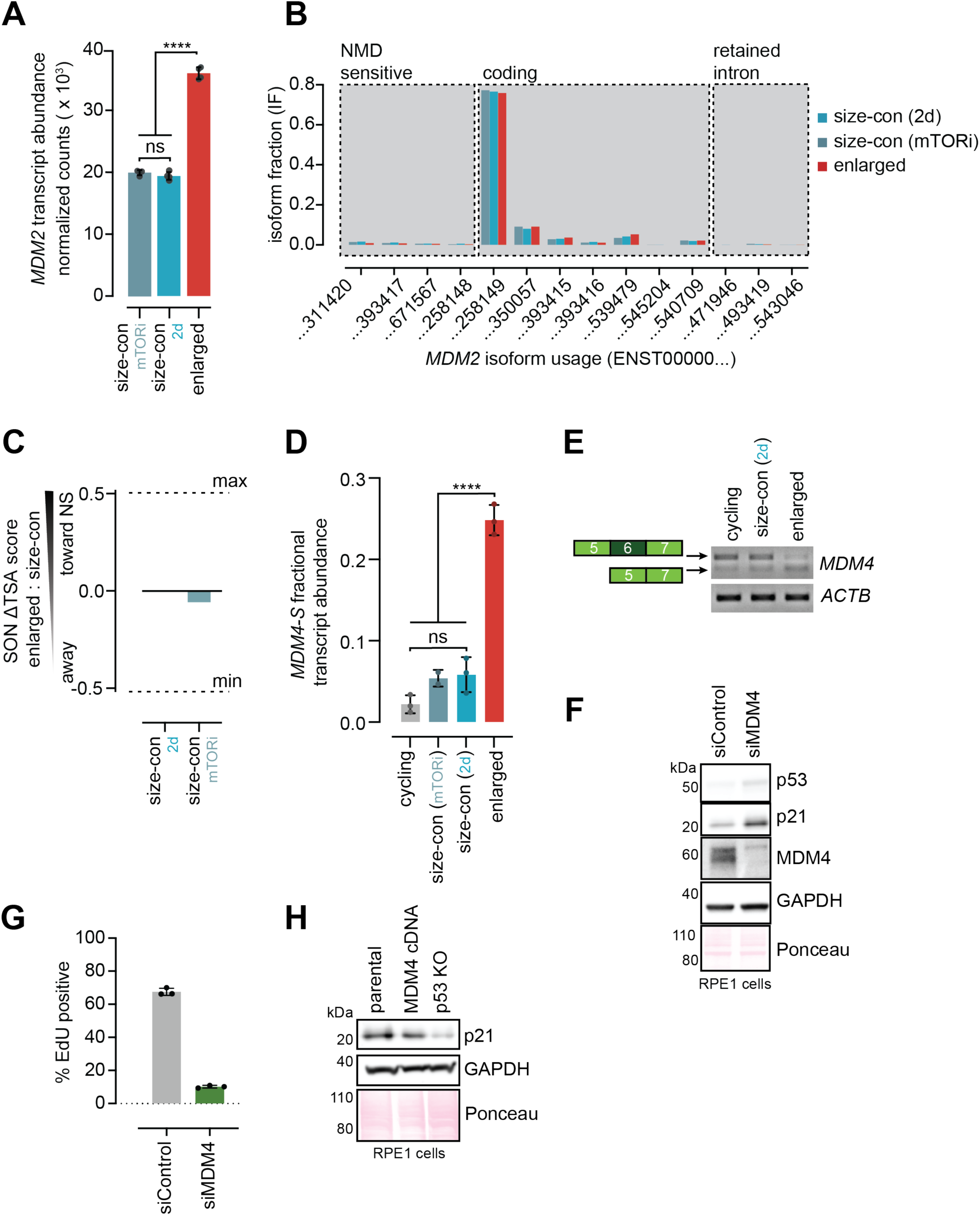
**A.** *MDM2* transcript abundance in enlarged and size-constrained cells as measured by RNAseq. Data are reported as mean ± SD from four replicates. ns: p = 0.48; **** : p < 0.0001 by one-way ANOVA. **B.** Isoform fractions for the detected isoforms of *MDM2* in the RNAseq dataset in enlarged and size-constrained cells. **C.** ΔTSA scores for *MDM4* in enlarged vs. size-constrained cells. Size-constraint method is indicated on the x-axis. Dashed lines indicate the maximum and minimum ΔTSA scores in the dataset. **D.** Fractional abundance measurements of the MDM4-S transcript variant by RT-qPCR in cycling, size-constrained, and enlarged cells. Error bars: mean ± SD for 2-3 biological replicates. ns : p ≥ 0.12; **** : p < 0.0001 by one-way ANOVA followed by Tukey’s multiple comparisons test. **E.** RT-PCR detecting *MDM4-FL* and *MDM4-S* in cycling, size-constrained (2d), and enlarged cells. ACTB was used as a loading control. **F.** Western blot depicting p53 and p21 protein levels in asynchronous RPE1 cells treated with an siRNA directed against MDM4. GAPDH and Ponceau staining are used as loading controls. **G.** Asynchronous RPE1 cells were transfected with the indicated siRNAs for 24 hours before being exposed to media containing EdU for an additional 24 hours. Cells were then fixed and stained for EdU. The fraction of EdU-positive cells is quantified for each condition in three technical replicates. **H.** Western blot depicting p21 protein levels in the indicated asynchronous RPE1 cell lines. GAPDH and Ponceau staining are used as loading controls.

**Figure S7: Related to Figure 6.**
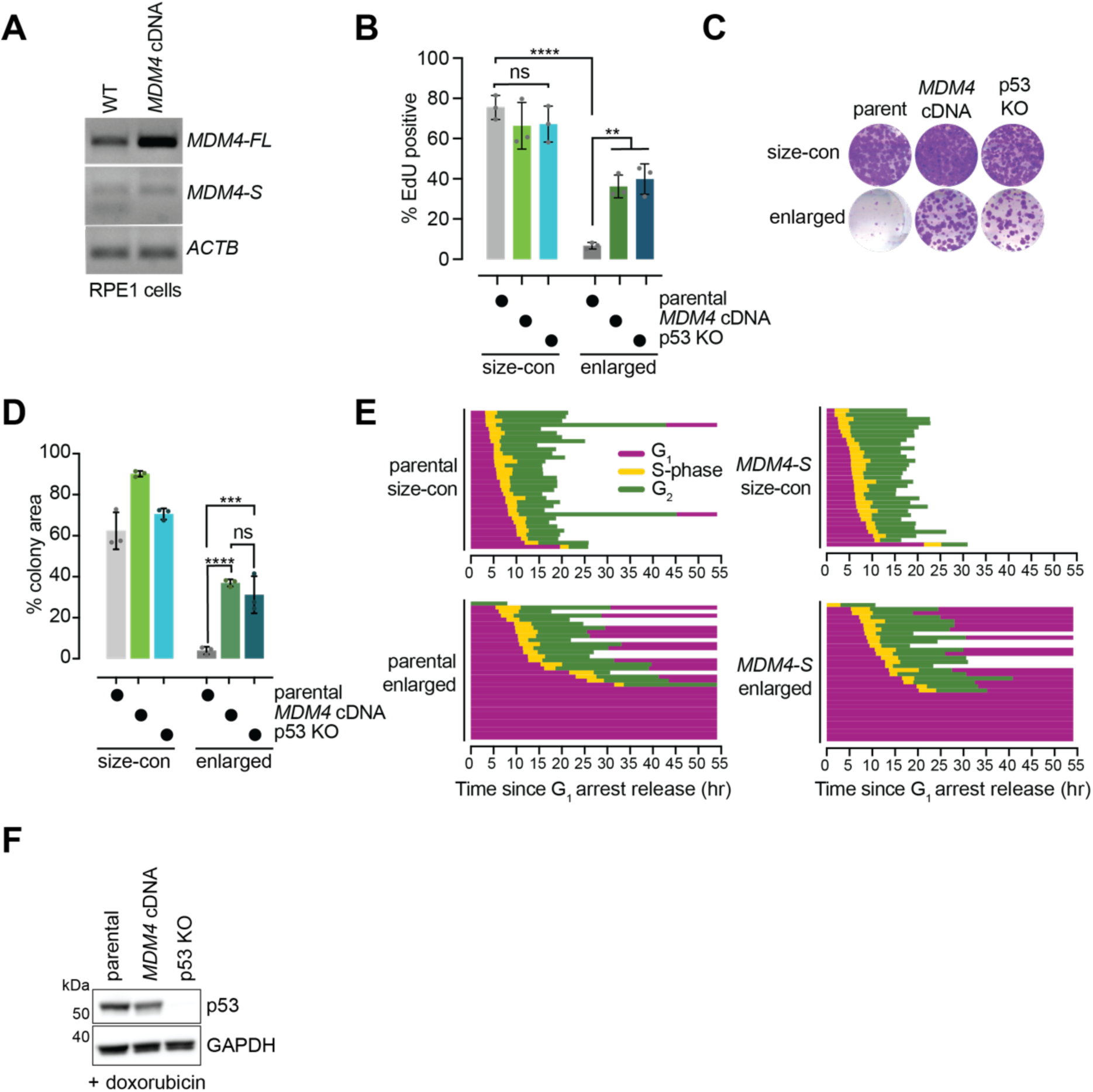
**A.** *MDM4-FL* and *MDM4-S* levels in parental and *MDM4* cDNA-expressing RPE1 cells by RT-PCR. **B.** The indicated stable RPE1 cell lines were treated as indicated and then washed into EdU-containing media for 24 hours. Cells were then fixed and stained for EdU. The fraction of EdU-positive cells is quantified for each condition. ns : p > 0.9; **** : p < 0.0001; ** : p < 0.007 by two-way ANOVA followed by Šidák’s multiple comparisons test. Error bars indicate mean ± SD for three replicates. **C.** Example wells for a colony formation assay where the indicated stable RPE1 cell lines released from the indicated G_1_ arrest and allowed to form colonies for 10 days. Colonies were stained with crystal violet. **D.** Quantification of the experiment shown in (**C**). ns: p = 0.53; *** : p = 0.0002; **** : p < 0.0001 by two-way ANOVA followed by Šidák’s multiple comparisons test. Error bars indicate mean ± SD for three replicates. **E.** Single cell traces for size-constrained (mTORi) and enlarged cells in the indicated RPE1 FUCCI stable cell lines released from a G_1_ arrest based on live cell imaging of FUCCI reporters. The end of a trajectory indicates mitotic entry or the end of the imaging duration. n= 34 cells per condition. **F.** Western blot depicting p53 levels in the indicated stable RPE1 cell lines treated with doxorubicin for 24 hours. GAPDH is used as a loading control.

## References

1. Manohar, S. and G.E. Neurohr, Too big not to fail: emerging evidence for size-induced senescence. FEBS J, 2024. 291(11): p. 2291–2305.

2. Dolznig, H., F. Grebien, T. Sauer, H. Beug, and E.W. Mullner, Evidence for a size-sensing mechanism in animal cells. Nat Cell Biol, 2004. 6(9): p. 899–905.

3. Ginzberg, M.B., N. Chang, H. D’Souza, N. Patel, R. Kafri, and M.W. Kirschner, Cell size sensing in animal cells coordinates anabolic growth rates and cell cycle progression to maintain cell size uniformity. eLife, 2018. 7: p. e26957.

4. Ginzberg, M.B., R. Kafri, and M. Kirschner, Cell biology. On being the right (cell) size. Science, 2015. 348(6236): p. 1245075.

5. Yilmaz, E., Z. Fan, J.P. Beech, V.S. Swaminathan, and J.O. Tegenfeldt, Size-based sorting of cancer cells reveals functional heterogeneity among subpopulations. Lab Chip, 2026. 26(11): p. 3459–3472.

6. Asadullah, S. Kumar, N. Saxena, M. Sarkar, A. Barai, and S. Sen, Combined heterogeneity in cell size and deformability promotes cancer invasiveness. J Cell Sci, 2021. 134(7).

7. Liu, S., M.B. Ginzberg, N. Patel, M. Hild, B. Leung, Z. Li, Y.C. Chen, N. Chang, Y. Wang, C. Tan, S. Diena, W. Trimble, L. Wasserman, J.L. Jenkins, M.W. Kirschner, and R. Kafri, Size uniformity of animal cells is actively maintained by a p38 MAPK-dependent regulation of G1-length. eLife, 2018. 7: p. e26947.

8. Hayflick, L. and P.S. Moorhead, The serial cultivation of human diploid cell strains. Exp Cell Res, 1961. 25(3): p. 585–621.

9. Manohar, S., M.E. Estrada, F. Uliana, K. Vuina, P.M. Alvarez, R.A.M. de Bruin, and G.E. Neurohr, Genome homeostasis defects drive enlarged cells into senescence. Mol Cell, 2023. 83(22): p. 4032–4046.

10. Crozier, L., R. Foy, R. Adib, A. Kar, J.A. Holt, A.U. Pareri, J.M. Valverde, R. Rivera, W.A. Weston, R. Wilson, C. Regnault, P. Whitfield, M. Badonyi, L.G. Bennett, E.G. Vernon, A. Gamble, J.A. Marsh, C.J. Staples, A.T. Saurin, A.R. Barr, and T. Ly, CDK4/6 inhibitor-mediated cell overgrowth triggers osmotic and replication stress to promote senescence. Mol Cell, 2023. 83(22): p. 4062–4077 e5.

11. Crozier, L., R. Foy, B.L. Mouery, R.H. Whitaker, A. Corno, C. Spanos, T. Ly, J. Gowen Cook, and A.T. Saurin, CDK4/6 inhibitors induce replication stress to cause long-term cell cycle withdrawal. EMBO J, 2022. 41(6): p. e108599.

12. Foy, R., L. Crozier, A.U. Pareri, J.M. Valverde, B.H. Park, T. Ly, and A.T. Saurin, Oncogenic signals prime cancer cells for toxic cell overgrowth during a G1 cell cycle arrest. Mol Cell, 2023. 83(22): p. 4047–4061.

13. Pareri, A.U., M. Losito, F. Foijer, and A.T. Saurin, Cell enlargement causes mitotic errors and aneuploidy in cells that evade senescence after CDK4/6 inhibition. bioRxiv, 2025: p. 2025.07.01.662622.

14. Neurohr, G.E., R.L. Terry, J. Lengefeld, M. Bonney, G.P. Brittingham, F. Moretto, T.P. Miettinen, L.P. Vaites, L.M. Soares, J.A. Paulo, J.W. Harper, S. Buratowski, S. Manalis, F.J. van Werven, L.J. Holt, and A. Amon, Excessive Cell Growth Causes Cytoplasm Dilution And Contributes to Senescence. Cell, 2019. 176(5): p. 1083–1097 e18.

15. Killander, D. and A. Zetterberg, A quantitative cytochemical investigation of the relationship between cell mass and initiation of DNA synthesis in mouse fibroblasts in vitro. Exp Cell Res, 1965. 40(1): p. 12–20.

16. Zatulovskiy, E., S. Zhang, D.F. Berenson, B.R. Topacio, and J.M. Skotheim, Cell growth dilutes the cell cycle inhibitor Rb to trigger cell division. Science, 2020. 369(6502): p. 466–471.

17. Xie, S., S. Zhang, G. de Medeiros, P. Liberali, and J.M. Skotheim, The G1/S transition in mammalian stem cells in vivo is autonomously regulated by cell size. Nat Commun, 2025. 16(1): p. 9071.

18. Fischer, M., Census and evaluation of p53 target genes. Oncogene, 2017. 36(28): p. 3943–3956.

19. Sakaue-Sawano, A., H. Kurokawa, T. Morimura, A. Hanyu, H. Hama, H. Osawa, S. Kashiwagi, K. Fukami, T. Miyata, H. Miyoshi, T. Imamura, M. Ogawa, H. Masai, and A. Miyawaki, Visualizing spatiotemporal dynamics of multicellular cell-cycle progression. Cell, 2008. 132(3): p. 487–98.

20. Krenning, L., Femke M. Feringa, Indra A. Shaltiel, J. van den Berg, and R.H. Medema, Transient Activation of p53 in G2 Phase Is Sufficient to Induce Senescence. Mol Cell, 2014. 55(1): p. 59–72.

21. Johmura, Y., M. Shimada, T. Misaki, A. Naiki-Ito, H. Miyoshi, N. Motoyama, N. Ohtani, E. Hara, M. Nakamura, A. Morita, S. Takahashi, and M. Nakanishi, Necessary and Sufficient Role for a Mitosis Skip in Senescence Induction. Mol Cell, 2014. 55(1): p. 73–84.

22. Wentinck, K., C. Li, N. Noureddine, A. Kitanovic, S. Manohar, F. Uliana, B. Meier, W. Zhao, E. Petsalaki, R.J. Morscher, G.E. Neurohr, and T. Kleele, GSK3β drives mitochondrial dysfunction and metabolic reprogramming in cellular senescence. bioRxiv, 2026.

23. Lanz, M.C., E. Zatulovskiy, M.P. Swaffer, L. Zhang, S. Zhang, D.S. You, G. Marinov, P. McAlpine, J.E. Elias, and J.M. Skotheim, Increasing cell size remodels the proteome and promotes senescence. Mol Cell, 2022: p. 2021.07.29.454227.

24. Karasu, M.E., L. Jahnke, B.J. Joseph, Y. Amerzhanova, A. Mironov, X. Shu, M.S. Schroder, A. Gvozdenovic, I. Sala, M. Zavolan, S. Jonas, and J.E. Corn, CCAR1 promotes DNA repair via alternative splicing. Mol Cell, 2024. 84(14): p. 2634–2647 e9.

25. Ilik, I.A. and T. Aktas, Nuclear speckles: dynamic hubs of gene expression regulation. FEBS J, 2022. 289(22): p. 7234–7245.

26. Zhang, M., Z. Gu, S. Guo, Y. Sun, S. Ma, S. Yang, J. Guo, C. Fang, L. Shu, Y. Ge, Z. Chen, and B. Wang, SRRM2 phase separation drives assembly of nuclear speckle subcompartments. Cell Rep, 2024. 43(3): p. 113827.

27. Ilik, I.A., M. Malszycki, A.K. Lubke, C. Schade, D. Meierhofer, and T. Aktas, SON and SRRM2 are essential for nuclear speckle formation. eLife, 2020. 9: p. e60579.

28. Fei, J.Y., M. Jadaliha, T.S. Harmon, I.T.S. Li, B.Y. Hua, Q.Y. Hao, A.S. Holehouse, M. Reyer, Q.Y. Sun, S.M. Freier, R.V. Pappu, K.V. Prasanth, and T. Ha, Quantitative analysis of multilayer organization of proteins and RNA in nuclear speckles at super resolution. J Cell Sci, 2017. 130(24): p. 4180–4192.

29. Kim, J., K.Y. Han, N. Khanna, T. Ha, and A.S. Belmont, Nuclear speckle fusion via long-range directional motion regulates speckle morphology after transcriptional inhibition. J Cell Sci, 2019. 132(8).

30. Wu, J., Y. Xiao, Y. Liu, L. Wen, C. Jin, S. Liu, S. Paul, C. He, O. Regev, and J. Fei, Dynamics of RNA localization to nuclear speckles are connected to splicing efficiency. Sci Adv, 2024. 10(42): p. eadp7727.

31. Neumann, F.R. and P. Nurse, Nuclear size control in fission yeast. J Cell Biol, 2007. 179(4): p. 593–600.

32. Jorgensen, P., N.P. Edgington, B.L. Schneider, I. Rupes, M. Tyers, and B. Futcher, The size of the nucleus increases as yeast cells grow. Mol Biol Cell, 2007. 18(9): p. 3523–32.

33. Chen, Y., Y. Zhang, Y. Wang, L. Zhang, E.K. Brinkman, S.A. Adam, R. Goldman, B. van Steensel, J. Ma, and A.S. Belmont, Mapping 3D genome organization relative to nuclear compartments using TSA-Seq as a cytological ruler. J Cell Biol, 2018. 217(11): p. 4025–4048.

34. Zhang, L., Y. Zhang, Y. Chen, O. Gholamalamdari, Y. Wang, J. Ma, and A.S. Belmont, TSA-seq reveals a largely conserved genome organization relative to nuclear speckles with small position changes tightly correlated with gene expression changes. Genome Res, 2020. 31(2): p. 251–264.

35. Kim, J., N.C. Venkata, G.A. Hernandez Gonzalez, N. Khanna, and A.S. Belmont Gene expression amplification by nuclear speckle association. J Cell Biol, 2019. 219(1).

36. Bhat, P., A. Chow, B. Emert, O. Ettlin, S.A. Quinodoz, M. Strehle, Y. Takei, A. Burr, I.N. Goronzy, A.W. Chen, W. Huang, J.L.M. Ferrer, E. Soehalim, S.T. Goh, T. Chari, D.K. Sullivan, M.R. Blanco, and M. Guttman, Genome organization around nuclear speckles drives mRNA splicing efficiency. Nature, 2024. 629(8014): p. 1165–1173.

37. Berry, S. and L. Pelkmans, Mechanisms of cellular mRNA transcript homeostasis. Trends Cell Biol, 2022. 32(8): p. 655–668.

38. Gholamalamdari, O., T. van Schaik, Y. Wang, P. Kumar, L. Zhang, Y. Zhang, G.A.H. Gonzalez, A.E. Vouzas, P.A. Zhao, D.M. Gilbert, J. Ma, B. van Steensel, and A.S. Belmont, Beyond A and B Compartments: how major nuclear locales define nuclear genome organization and function. eLife, 2025.

39. Wang, Y., Y. Zhang, R. Zhang, T. van Schaik, L. Zhang, T. Sasaki, D. Peric-Hupkes, Y. Chen, D.M. Gilbert, B. van Steensel, A.S. Belmont, and J. Ma, SPIN reveals genome-wide landscape of nuclear compartmentalization. Genome Biology, 2021. 22(1): p. 36.

40. Tran, J.R., X. Zheng, S.A. Adam, R.D. Goldman, and Y. Zheng, High quality mapping of chromatin at or near the nuclear lamina from small numbers of cells reveals cell cycle and developmental changes of chromatin at the nuclear periphery. Nucleic Acids Res, 2022. 50(20): p. e117.

41. Namba, H., S. Mir, E.L. Groce, and K.A. Alexander, Nuclear speckles: a fundamental layer of gene regulation. Trends in Cell Biol, 2026.

42. Guelen, L., L. Pagie, E. Brasset, W. Meuleman, M.B. Faza, W. Talhout, B.H. Eussen, A. de Klein, L. Wessels, W. de Laat, and B. van Steensel, Domain organization of human chromosomes revealed by mapping of nuclear lamina interactions. Nature, 2008. 453(7197): p. 948–51.

43. Kashirina, D.N., A.Y. Ratushny, D.K. Matveeva, M.I. Ezdakova, L.K. Pastushkova, I.M. Larina, and L.B. Buravkova, Modifications of the Secretome of Mesenchymal Stromal Cells under Conditions of Stress-Induced Aging. Bull Exp Biol Med, 2025. 179(4): p. 518–525.

44. Coppe, J.P., P.Y. Desprez, A. Krtolica, and J. Campisi, The senescence-associated secretory phenotype: the dark side of tumor suppression. Annu Rev Pathol, 2010. 5: p. 99–118.

45. Park, J.J., S.J. Lee, M. Baek, O.J. Lee, S. Nam, J. Kim, J.Y. Kim, E.Y. Shin, and E.G. Kim, FRMD6 determines the cell fate towards senescence: involvement of the Hippo-YAP-CCN3 axis. Cell Death Differ, 2024. 31(11): p. 1398–1409.

46. Feng, D., J. Wang, D. Li, R. Wu, W. Wei, and C. Zhang, Senescence-associated secretory phenotype constructed detrimental and beneficial subtypes and prognostic index for prostate cancer patients undergoing radical prostatectomy. Discov Oncol, 2023. 14(1): p. 155.

47. Basisty, N., A. Kale, O.H. Jeon, C. Kuehnemann, T. Payne, C. Rao, A. Holtz, S. Shah, V. Sharma, L. Ferrucci, J. Campisi, and B. Schilling, A proteomic atlas of senescence-associated secretomes for aging biomarker development. PLoS Biol, 2020. 18(1): p. e3000599.

48. Birch, J. and J. Gil, Senescence and the SASP: many therapeutic avenues. Genes Dev, 2020. 34(23-24): p. 1565–1576.

49. Lehrer, S. and P. Rheinstein, CDKN2A mRNA Over-expression Is Associated With CD8+ T Cell Exclusion and IDO1-Mediated Adaptive Immune Resistance in Gastric Adenocarcinoma. Cancer Diagn Progn, 2026. 6(4): p. 639–648.

50. Acosta, J.C., A. Banito, T. Wuestefeld, A. Georgilis, P. Janich, J.P. Morton, D. Athineos, T.W. Kang, F. Lasitschka, M. Andrulis, G. Pascual, K.J. Morris, S. Khan, H. Jin, G. Dharmalingam, A.P. Snijders, T. Carroll, D. Capper, C. Pritchard, G.J. Inman, T. Longerich, O.J. Sansom, S.A. Benitah, L. Zender, and J. Gil, A complex secretory program orchestrated by the inflammasome controls paracrine senescence. Nat Cell Biol, 2013. 15(8): p. 978–90.

51. Marchal, C., J. Sima, and D.M. Gilbert, Control of DNA replication timing in the 3D genome. Nat Rev Mol Cell Biol, 2019. 20(12): p. 721–737.

52. Orengo, J.P., D. Bundman, and T.A. Cooper, A bichromatic fluorescent reporter for cell-based screens of alternative splicing. Nucleic Acids Res, 2006. 34(22): p. e148.

53. Liang, Y., T. Tebaldi, K. Rejeski, P. Joshi, G. Stefani, A. Taylor, Y. Song, R. Vasic, J. Maziarz, K. Balasubramanian, A. Ardasheva, A. Ding, A. Quattrone, and S. Halene, SRSF2 mutations drive oncogenesis by activating a global program of aberrant alternative splicing in hematopoietic cells. Leukemia, 2018. 32(12): p. 2659–2671.

54. Yano, K., R.U. Takahashi, B. Shiotani, J. Abe, T. Shidooka, Y. Sudo, Y. Yamamoto, S. Kan, H. Sakagami, and H. Tahara, PRPF19 regulates p53-dependent cellular senescence by modulating alternative splicing of MDM4 mRNA. J Biol Chem, 2021. 297(1): p. 100882.

55. Deschenes, M., M. Durand, M.A. Olivier, A. Pellerin-Viger, F. Rodier, and B. Chabot, A defective splicing machinery promotes senescence through MDM4 alternative splicing. Aging Cell, 2024. n/a(n/a): p. e14301.

56. Beauchamp, M.-C., A. Djedid, E. Bareke, F. Merkuri, R. Aber, A.S. Tam, M.A. Lines, K.M. Boycott, P.C. Stirling, J.L. Fish, J. Majewski, and L.A. Jerome-Majewska, Mutation in Eftud2 causes craniofacial defects in mice via mis-splicing of Mdm2 and increased P53. Hum Mol Genet, 2021. 30: p. 739–757.

57. Kwon, S.M., S. Min, U.-w. Jeoun, M.S. Sim, G.H. Jung, S.M. Hong, B.A. Jee, H.G. Woo, C. Lee, and G. Yoon, Global spliceosome activity regulates entry into cellular senescence. FASEB J, 2021. 35(1): p. e21204.

58. Deschenes, M. and B. Chabot, The emerging role of alternative splicing in senescence and aging. Aging Cell, 2017. 16(5): p. 918–933.

59. Barak, Y., T. Juven, R. Haffner, and M. Oren, mdm2 expression is induced by wild type p53 activity. EMBO J, 1993. 12(2): p. 461–8.

60. Wu, X., J.H. Bayle, D. Olson, and A.J. Levine, The p53-mdm-2 autoregulatory feedback loop. Genes Dev, 1993. 7(7A): p. 1126–32.

61. Criscione, S.W., Y.V. Teo, and N. Neretti, The Chromatin Landscape of Cellular Senescence. Trends Genet, 2016. 32(11): p. 751–761.

62. Yu, R., H. Sas-Nowosielska, A.S. Dessai, A. Oubarri, H. Kim, C.L. Faunce, B. Vanderkruk, C.R. Good, K.A. Alexander, S.C. Nguyen, I.D. Krantz, E.F. Joyce, M. Lakadamyali, and S.L. Berger, SWING regions prime chromatin for nuclear speckle–mediated gene regulation. bioRxiv, 2026: p. 2025.10.01.679801.

63. Ahanger, S.H., E.R. Semenza, C. Zhang, E. Gil, M.A. Cole, S.H. Lu, L. Wang, A.R. Kriegstein, and D.A. Lim, Subnuclear genome compartmentalization controls bivalent chromatin activity. Nature, 2026. 657(8131): p. 539–548.

64. Solovei, I., A.S. Wang, K. Thanisch, C.S. Schmidt, S. Krebs, M. Zwerger, T.V. Cohen, D. Devys, R. Foisner, L. Peichl, H. Herrmann, H. Blum, D. Engelkamp, C.L. Stewart, H. Leonhardt, and B. Joffe, LBR and lamin A/C sequentially tether peripheral heterochromatin and inversely regulate differentiation. Cell, 2013. 152(3): p. 584–98.

65. Peric-Hupkes, D., W. Meuleman, L. Pagie, S.W. Bruggeman, I. Solovei, W. Brugman, S. Graf, P. Flicek, R.M. Kerkhoven, M. van Lohuizen, M. Reinders, L. Wessels, and B. van Steensel, Molecular maps of the reorganization of genome-nuclear lamina interactions during differentiation. Mol Cell, 2010. 38(4): p. 603–13.

66. Balachandra, S., S. Sarkar, and A.A. Amodeo, The Nuclear-to-Cytoplasmic Ratio: Coupling DNA Content to Cell Size, Cell Cycle, and Biosynthetic Capacity. Annu Rev Genet, 2022. 56: p. 165–185.

67. Marchingo, J.M., L.V. Sinclair, A.J. Howden, and D.A. Cantrell, Quantitative analysis of how Myc controls T cell proteomes and metabolic pathways during T cell activation. eLife, 2020. 9: p. e53725.

68. Waugh, R.E., E. Lomakina, A. Amitrano, and M. Kim, Activation effects on the physical characteristics of T lymphocytes. Front Bioeng Biotechnol, 2023. 11: p. 1175570.

69. Palikyras, S., V. Varamogiani-Mamatsi, Y. Zhu, A. Mizi, S. Ramasamy, J.V. Grini, I. Liebermann, A. Stavropoulou, A.P. Puig, T.S. Ogurlu-Tank, K. Kydonakis, I. Papadionysiou, D. Bartsch, Y. Kargapolova, N.R. Morero, K. Sofiadis, C. Nikolaou, F. Kundel, L. Kurian, J. Ellenberg, A.M. Oudelaar, M. Barbieri, K.S. Beckwith, and A. Papantonis, Senescent cells cluster CTCF on nuclear speckles to instruct an alternative splicing program. Nat Aging, 2026. 6(8): p. 1592–1610.

70. Kim, J., Y. Lee, T. Jeon, S. Ju, J.S. Kim, M.S. Kim, and C. Kang, Autophagy-dependent splicing control directs translation toward inflammation during senescence. Dev Cell, 2025. 60(3): p. 364–378 e7.

71. AbuHammad, S., C. Cullinane, C. Martin, Z. Bacolas, T. Ward, H. Chen, A. Slater, K. Ardley, L. Kirby, K.T. Chan, N. Brajanovski, L.K. Smith, A.D. Rao, E.J. Lelliott, M. Kleinschmidt, I.A. Vergara, A.T. Papenfuss, P. Lau, P. Ghosh, S. Haupt, Y. Haupt, E. Sanij, G. Poortinga, R.B. Pearson, H. Falk, D.J. Curtis, P. Stupple, M. Devlin, I. Street, M.A. Davies, G.A. McArthur, and K.E. Sheppard, Regulation of PRMT5-MDM4 axis is critical in the response to CDK4/6 inhibitors in melanoma. Proc Natl Acad Sci U S A, 2019. 116(36): p. 17990–18000.

72. Soliman, M.A., P. Berardi, S. Pastyryeva, P. Bonnefin, X. Feng, A. Colina, D. Young, and K. Riabowol, ING1a expression increases during replicative senescence and induces a senescent phenotype. Aging Cell, 2008. 7(6): p. 783–94.

73. Brueckner, L., P.A. Zhao, T. van Schaik, C. Leemans, J. Sima, D. Peric-Hupkes, D.M. Gilbert, and B. van Steensel, Local rewiring of genome-nuclear lamina interactions by transcription. EMBO J, 2020. 39(6): p. e103159.

74. Gibson, D.G., L. Young, R.Y. Chuang, J.C. Venter, C.A. Hutchison, 3rd, and H.O. Smith, Enzymatic assembly of DNA molecules up to several hundred kilobases. Nat Methods, 2009. 6(5): p. 343–5.

75. Manohar, S., Q. Yu, S.P. Gygi, and R.W. King, The Insulin Receptor Adaptor IRS2 is an APC/C Substrate That Promotes Cell Cycle Protein Expression and a Robust Spindle Assembly Checkpoint. Mol Cell Proteomics, 2020. 19(9): p. 1450–1467.

76. Chen, S., Y. Zhou, Y. Chen, and J. Gu, fastp: an ultra-fast all-in-one FASTQ preprocessor. Bioinformatics, 2018. 34(17): p. i884–i890.

77. Bray, N.L., H. Pimentel, P. Melsted, and L. Pachter, Near-optimal probabilistic RNA-seq quantification. Nat Biotechnol, 2016. 34(5): p. 525–7.

78. Vitting-Seerup, K. and A. Sandelin, IsoformSwitchAnalyzeR: analysis of changes in genome-wide patterns of alternative splicing and its functional consequences. Bioinformatics, 2019. 35(21): p. 4469–4471.

79. Langmead, B. and S.L. Salzberg, Fast gapped-read alignment with Bowtie 2. Nat Methods, 2012. 9(4): p. 357–9.

80. Dekker, J., A.S. Belmont, M. Guttman, V.O. Leshyk, J.T. Lis, S. Lomvardas, L.A. Mirny, C.C. O’Shea, P.J. Park, B. Ren, J.C.R. Politz, J. Shendure, S. Zhong, and D.N. Network, The 4D nucleome project. Nature, 2017. 549(7671): p. 219–226.

81. Reiff, S.B., A.J. Schroeder, K. Kirli, A. Cosolo, C. Bakker, L. Mercado, S. Lee, A.D. Veit, A.K. Balashov, C. Vitzthum, W. Ronchetti, K.M. Pitman, J. Johnson, S.R. Ehmsen, P. Kerpedjiev, N. Abdennur, M. Imakaev, S.U. Ozturk, U. Camoglu, L.A. Mirny, N. Gehlenborg, B.H. Alver, and P.J. Park, The 4D Nucleome Data Portal as a resource for searching and visualizing curated nucleomics data. Nat Commun, 2022. 13(1): p. 2365.

82. Schindelin, J., I. Arganda-Carreras, E. Frise, V. Kaynig, M. Longair, T. Pietzsch, S. Preibisch, C. Rueden, S. Saalfeld, B. Schmid, J.Y. Tinevez, D.J. White, V. Hartenstein, K. Eliceiri, P. Tomancak, and A. Cardona, Fiji: an open-source platform for biological-image analysis. Nat Methods, 2012. 9(7): p. 676–82.

83. Jansen, J., K.E. Bohnsack, S. Bohlken-Fascher, M.T. Bohnsack, and M. Dobbelstein, The ribosomal protein L22 binds the MDM4 pre-mRNA and promotes exon skipping to activate p53 upon nucleolar stress. Cell Rep, 2024. 43(8): p. 114610.

84. Guzman, C., M. Bagga, A. Kaur, J. Westermarck, and D. Abankwa, ColonyArea: an ImageJ plugin to automatically quantify colony formation in clonogenic assays. PLoS One, 2014. 9(3): p. e92444.

85. Ge, S.X., D. Jung, and R. Yao, ShinyGO: a graphical gene-set enrichment tool for animals and plants. Bioinformatics, 2020. 36(8): p. 2628–2629.

